# A spatial map of human macrophage niches links tissue location with function

**DOI:** 10.1101/2022.08.18.504434

**Authors:** Magdalena Matusiak, John W. Hickey, Bogdan Luca, Guolan Lu, Lukasz Kidzinski, Shirley Shu, Deana Rae Crystal Colburg, Darci J. Phillips, Sky W. Brubaker, Gregory W. Charville, Jeanne Shen, Garry P. Nolan, Aaron M. Newman, Robert B. West, Matt van de Rijn

## Abstract

Macrophages are the most abundant immune cell type in the tumor microenvironment (TME). Yet the spatial distribution and cell interactions that shape macrophage function are incompletely understood. Here we use single-cell RNA sequencing data and multiplex imaging to discriminate and spatially resolve macrophage niches within benign and malignant breast and colon tissue. We discover four distinct tissue-resident macrophage (TRM) layers within benign bowel, two TRM niches within benign breast, and three tumor-associated macrophage (TAM) populations within breast and colon cancer. We demonstrate that IL4I1 marks phagocytosing macrophages, SPP1 TAMs are enriched in hypoxic and necrotic tumor regions, and a novel subset of FOLR2 TRMs localizes within the plasma cell niche. Furthermore, NLRP3 TAMs that colocalize with neutrophils activate an inflammasome in the TME and in Crohn’s disease and are associated with poor outcomes in breast cancer patients. This work suggests novel macrophage therapy targets and provides a framework to study human macrophage function in clinical samples.

## Introduction

Tumor-associated macrophage (TAM) infiltration, as measured by CD68 macrophage infiltration, predicts poor patient outcomes for most tumor types (Fridman et al. 2017). This fact illustrates the critical role macrophages play in the TME. As a result, TAMs were surmised to be a promising cancer therapy target. However, TAM targeting therapeutic efforts have shown minimal single-agent efficacy against solid tumors, including CSF1 pathway blockade (Papadopoulos et al. 2017; Ries et al. 2014). This may be in part because such therapies treat macrophages as one entity and aim to repress macrophage biology broadly. Clearly, a better understanding of the molecular and functional diversity of TAMs is needed to facilitate rational macrophage targeting in cancer and predict clinical outcomes.

Previous studies revealed transcriptional macrophage heterogeneity in human cancer (Azizi et al. 2018; Zhang et al. 2020; Mulder et al. 2021). In addition, using immunostaining we and others showed that macrophage markers including MARCO, APOE, CCR2, TREM2, and FOLR2 are restricted to spatially discrete macrophage populations (La Fleur et al. 2018; Luca et al. 2021; Ramos et al. 2022) and demonstrated their differential spatial co-enrichment with distinct T cell subtypes (Luca et al. 2021; Ramos et al. 2022). However, these immunostaining studies were limited to examining one or two macrophage and T cell populations at once in one organ system. Unbiased and highly multiplexed profiling across different organ systems is needed to fully dissect macrophage spatial tissue organization and cell-cell interactions that shape macrophage functions in the TME.

Here we link single-cell RNA sequencing (scRNA Seq) data with multiplex immunofluorescence (IF) to discriminate five discrete macrophage populations in human tissues: benign breast and colon, breast cancer (BC), and colorectal cancer (CRC). By applying a cell co-enrichment analysis across all cell types, we reveal that the different macrophage populations occupy spatially distinct niches, which are characterized by unique cellular compositions and discrete functional properties. Specifically, we uncover a layered TRMs architecture in the benign colon and breast tissues. We then expand these findings to cancer and inflammatory diseases, showing that IL4I1 marks actively phagocytosing cells, a novel subset of FOLR2 TRMs is enriched in the plasma cell niche, SPP1 TAMs are associated with hypoxia and tumor necrosis, and NLRP3 TAMs activate the inflammasome in breast cancer (BC), colorectal cancer (CRC) and Crohn’s Disease (CD). Furthermore, we establish that NLRP3 TAMs with activated inflammasomes are spatially associated with neutrophil infiltration. Finally, NLRP3 TAMs and neutrophil niche abundance correlate with outcomes in BC patients and thus suggest NLRP3 inflammasome blockade as a novel therapeutic target in cancer and CD. This work conceptualizes the macrophage niche as a fundamental and conserved functional tissue building block and demonstrates strategies to identify and further study distinct macrophage populations *in situ* in human clinical specimens.

## Results

### Experimental approach

This work aimed to reveal the macrophage niches’ cellular composition and spatial tissue distribution. We chose to focus on BC and CRC because CD68 infiltration predicts BC and CRC patients outcome (Fridman et al. 2017; Beck et al. 2009). We first used four public scRNA Seq datasets of CRC and BC (Lee et al. 2020; Qian et al. 2020; Bassez et al. 2021) to discover markers of distinct macrophage subtypes **(Fig 1A**, results in **Fig1)**. Subsequently, we established a panel of 6 formalin-fixed, paraffin-embedded (FFPE)-compatible antibodies targeting transcriptional macrophage markers and identified five discrete macrophage populations *in situ*. Next, we used whole section immunohistochemistry (IHC), 4-color immunofluorescence (IF), and 36-antibody CODEX assays on Tissue Microarrays (TMAs) to discover distinct spatial macrophage niches and the possible functions these spatially-resolved macrophages fulfill in normal tissue and within the TME **(Fig 1B**, results in **Fig2-7)**.

**Fig 1.**
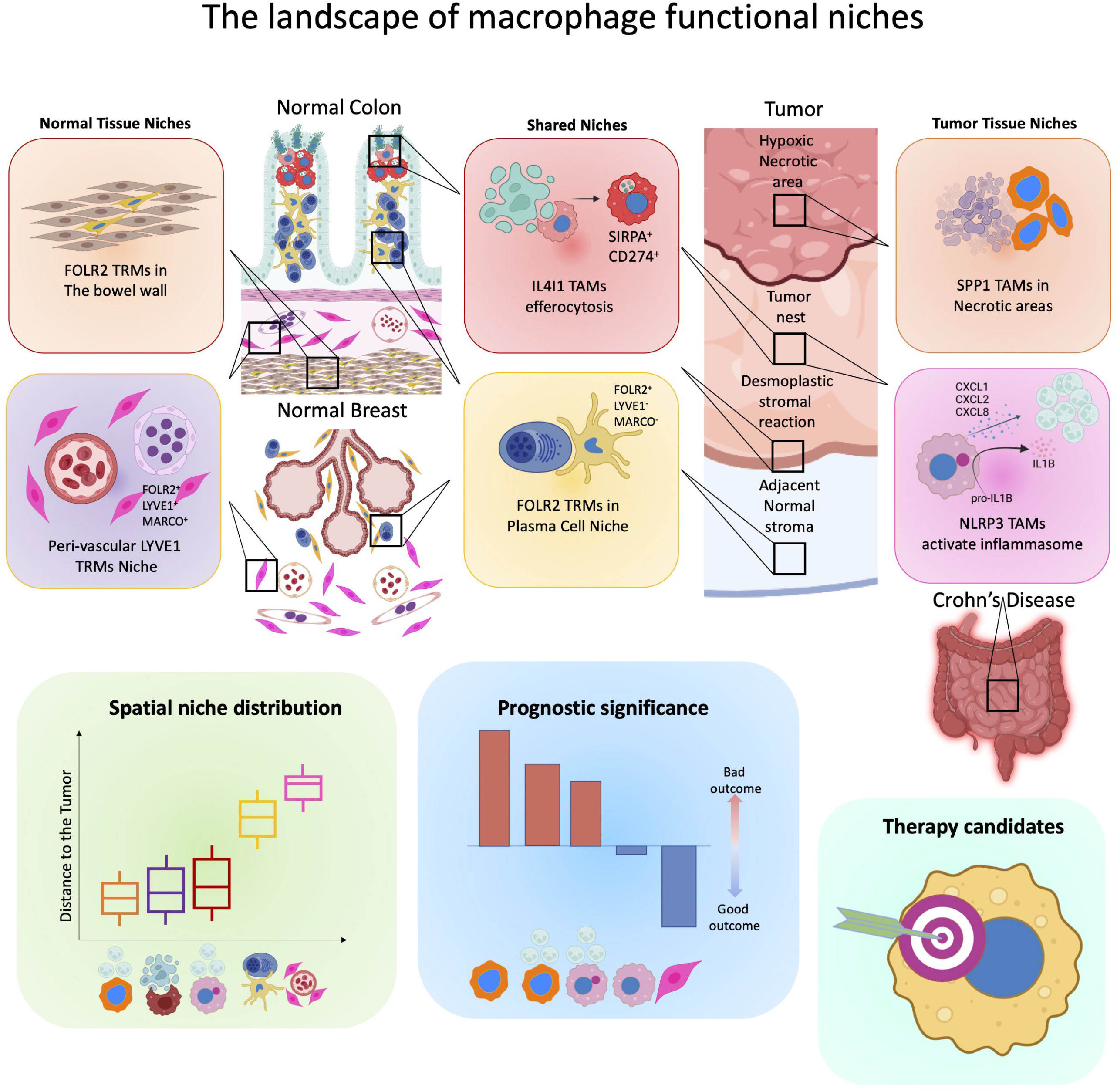
Single-cell RNA Sequencing and spatially resolved transcriptomics reveal differences in spatial enrichment of myeloid markers. (**A** and **B**) Flow charts of experimental design. (**C**) UMAP projection of monocyte and macrophage scRNA transcriptomes from 4 studies colored by annotated populations (*top*) and a breakdown of cells, samples and patient numbers by study (*bottom*). (**D**) Dotplot of average marker gene expression per scRNA mlyeloid populations. Highlighted in bold are 6 markers for which FFPE-compatible antibodies were identified. (**E**) Barplot of the ratio of log2 average fractional scRNA mmyeloid population enrichment between CRC and BC in tumor samples with more than 35 monocytes and macrophages detected. (**F**) Same as (**C**) but colored by anatomical location. (**G**) Same as (**E**) but between Normal colon samples and CRC samples in 2 CRC scRNA Seq datasets (Lee et al. 2020; Qian et al. 2020). (**H**) Immunofluorescence images show overlap of the established FFPE antibodies and CD68.

**Fig 2.**
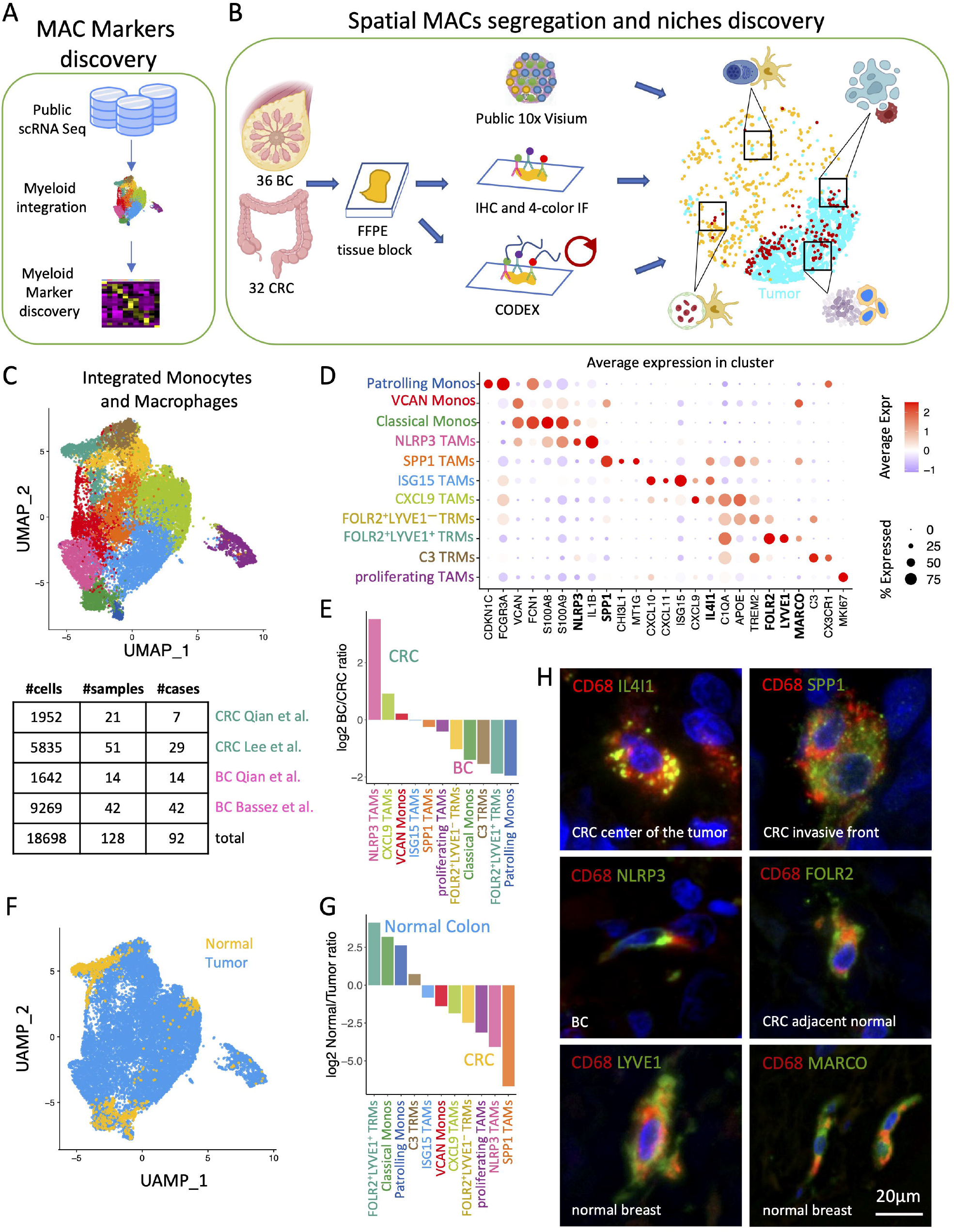
IL4I1, FOLR2, LYVE1, and MARCO label spatially segregated TRM niches in normal Colon and Breast. (**A**) Immunofluorescence (IF) images show 3 TRM layers marked by IL4I1, FOLR2, and LYVE1 in normal colon mucosa and submucosa. Note that LYVE1 also stains normal lymph vessels. (**B**) IF image shows that FOLR2^+^, LYVE1^+^ TRMs in normal colon submucosa are MARCO^+^. (**C**) Dotplot shows average FOLR2, LYVE1, and MARCO expression in scRNA macrophage populations. (**D**) Volcano plot shows top differentially expressed genes between FOLR2^+^, LYVE1^−^ and FOLR2^+^, LYVE1^+^ TRMs. (**E**) IF images show TRMs in normal breast marked by FOLR2, LYVE1, and MARCO, depending on whether they are Lobular (i) or Peri-vascular (ii). (**F**) The schematic shows the distribution of TRM populations colon mucosa and submucosa (*top*) and around normal breast glands (*bottom*). (**A**,**B**,**E**) Close-up images on the right correspond to boxed regions on the left. The scale bar of 10 μm is identical for all close-up images.

### ScRNA Seq and spatially resolved transcriptomics reveal differences in spatial enrichment of myeloid markers

To find markers for different macrophage populations, we integrated, clustered, and compared scRNA monocyte and macrophage transcriptomes from 18,698 cells from 92 patients across four published studies of BC and CRC (**Fig 1A, C, S1A**). We defined 11 transcriptional clusters marked by differential enrichment of genes (**Fig 1C-D**). We selected clustering resolution that separated known myeloid subtypes as follows: TRMs (*LYVE1*^*+*^) form TAMs (*TREM2*^*+*^*APOE*^*+*^), and Patrolling (*CDKN1C*^*+*^*FCGR3A*^*+*^) from Classical monocytes (*VCAN*^*+*^*S100A8*^*+*^*S100A9*^*+*^). We differentiated three monocyte, five TAM, and three TRM scRNA subsets and annotated them by their most differentially expressed genes. In agreement with previous reports (Qian et al. 2020; Mulder et al. 2021), we differentiated NLRP3 TAMs (*NLRP3*^*+*^ *IL1B*^*+*^), SPP1 TAMs (*SPP1*^*+*^*CHI3L1*^*+*^*MT1G*^*+*^), CXCL9 TAMs (*CXCL9*^*+*^*IL4I1*^*+*^), and ISG15 TAMs (*ISG15*^*+*^*CXCL10*^*+*^*CXCL11*^*+*^). In addition, we found three TRMs subsets: LYVE1^*-*^FOLR2^*+*^ (*FOLR2*^*+*^*APOE*^*+*^*TREM2*^*+*^), LYVE1^*+*^FOLR2^*+*^ TRMs (*FOLR2*^*+*^*LYVE1*^*+*^*MARCO*^*+*^*SLC40A1*^*+*^*SEPP1*^*+*^) and C3 TRMs (*C3*^*+*^*CX3CR1*^*+*^) (**Fig 1D**).

To explore the distribution of these subsets between CRC and BC, we computed a ratio of their average frequency across samples with more than 35 myeloid cells. CXCL9 TAMs were the most abundant TAM population in both BC and CRC and NLRP3 TAMs were enriched in CRC, with about 3.5 log2 fold higher frequency than in BC (**Fig 1E, S1B-D**). Next, we leveraged the fact that the two CRC datasets used (Qian et. al and Lee et. al) contained benign colon samples and compared macrophage cluster distribution across benign and tumor samples. We observed fundamental cluster segregation between benign colon and tumor tissue: LYVE1 TRMs were most enriched in benign colon, whereas NLRP3 TAMs and SPP1 TAMs were almost exclusively confined to colon tumors (**Fig 1F-G, S1E**). Next, guided by the differential marker gene enrichment between the 11 scRNA Seq myeloid subsets and the differences in their fractional enrichment between normal colon and CRC (**Fig 1D-G, S1E**), we built a panel of FFPE-compatible antibodies for six macrophage markers to resolve TAM and TRM populations (**Fig 1H**), including IL4I1, NLRP3, SPP1, FOLR2, LYVE1, and MARCO. The following sections describe how we used these markers to discriminate spatial macrophage niches (**Fig 2–3**) and to define their cellular composition and function (**Fig 4-7**).

### IL4I1, FOLR2, LYVE1, and MARCO label spatially segregated TRM niches in benign colon and breast

To comprehensively map the TRMs in benign human colon, we compared the protein expression of the canonical macrophage markers– CD68 and CD163– to our newly established macrophage marker panel. Interestingly, we found that TRMs near the surface (luminal aspect) of the lamina propria expressed more CD68 than CD163 (**Fig S2A i**), while the TRMs in the colon submucosa had higher CD163 compared to CD68 expression (**Fig S2A iii**). In contrast, TRMs inside the germinal center of the gut lymphoid follicles exhibited exceptionally high CD68 expression with a dim CD163 signal (**Fig S2A ii**). We further resolved the observed CD68-CD163 spatial differences using our newly established marker panel. Specifically, we revealed that IL4I1, FOLR2, LYVE1, and MARCO protein expression differentiated three distinct layers of TRMs in benign colon lamina propria (LP) and colon submucosa. IL4I1 marked TRMs along the colon crypt’s top (luminal aspect), while FOLR2 marked TRMs towards the middle and bottom of the crypt (**Fig 2A**). The third TRM layer was localized in the colon submucosa and was marked by FOLR2, LYVE1, and MARCO (**Fig 2B**). Since the gastrointestinal submucosa is rich in blood and lymph vessels, we termed the submucosal macrophage population peri-vascular (PV) FOLR2^+^LYVE1^+^MARCO^+^ TRMs. Of note, LYVE1 is also expressed on lymphatic endothelial cells, yet they can be readily differentiated from TRMs as they are organized in tubes, display much higher LYVE1 expression compared to TRMs, and do not express FOLR2 and MARCO (**Fig S4A**).

Furthermore, we showed that this TRM zonation is conserved across the intestinal tract, demonstrating that the three TRM layers were present in the duodenum (**Fig S2B**) and the ileum (**Fig S2C**). In addition to the differential protein marker expression, the three TRMs showed distinct morphological appearances; the IL4I1 TRMs at the top of LP had a foamy morphology, while the FOLR2 TRMs and LYVE1 TRMs displayed a spindle cell shape.

The striking spatial segregation of TRMs in the bowel wall is consistent with the scRNA Seq data that indicated the existence of two distinct FOLR2 TRMs populations: one that is positive for *FOLR2* alone (FOLR2^+^LYVE1^−^) and one that co-expresses *FOLR2, LYVE1*, and *MARCO* (FOLR2^+^LYVE1^+^, **Fig 2C**) as well as high levels of *CD163*. Differential gene expression showed that FOLR2^+^LYVE1^+^ TRMs are enriched in scavenger receptors (*MARCO, CD36, MRC1*), metabolic enzymes (*BLVRB, PDK4*), and immunoglobulins (*IGHA1, IGKC, IGLC2*). On the other hand, the FOLR2^+^LYVE1^−^ subset is enriched in phagocytosis and antigen presentation gene signatures, further supporting the distinct phenotypes of the two FOLR2 positive populations (**Fig 2D**).

Next, we examined TRM populations in benign breast stroma. Consistent with a recent report (Ramos et al. 2022), we show that TRMs surrounding benign breast lobules and ducts were FOLR2 positive (**Fig 2E**). We called these cells Lobular TRMs and found they express a dim level of LYVE1 and MARCO (**Fig 2E i**). Furthermore, we discovered that TRMs localized in the highly vascularized connective tissue that is further removed from the breast lobules co-expressed high levels of FOLR2, LYVE1, and MARCO (**Fig 2E ii**). The peri-vascular location and the same protein marker expression of the FOLR2^hi^LYVE1^hi^MARCO^hi^ TRMs from colon submucosa and breast connective tissue suggest they were the same cell population that is conserved across different organ systems. We did not detect any IL4I1 positive macrophages in the benign breast stroma (data not shown).

Our work significantly extends upon prior reports that found Lyve1 expression discriminated between murine TRM populations (Chakarov et al. 2019) and identified FOLR2 TRMs as humans’ main breast gland TRM population (Ramos et al. 2022). Specifically, we show that 1) two distinct FOLR2 positive TRM populations existed in human benign breast and colon, 2) that the LYVE1^hi^FOLR2^hi^ population also exhibited high MARCO expression, and 3) by providing a detailed topography of their distribution in benign colon and breast, and 4) by showing they were conserved across the breast and the GI tract (**Fig 2F**).

### FOLR2, IL4I1, NLRP3, and SPP1 mark spatially distinct macrophage niches in the TME

To study the spatial distribution of macrophage markers in the TME in breast and colon cancer, we used CD68 and CD163 as canonical macrophage markers, IL4I1, NLRP3, and SPP1 to differentiate scRNA TAM subsets, and FOLR2 to highlight TRMs. ScRNA Seq data indicated that NLRP3 is a specific NLRP3 TAM marker, SPP1 is a specific SPP1 TAM marker, and IL4I1 has a broader expression, highlighting SPP1 TAMs, CXCL9 TAMs, and ISG15 TAMs. Nevertheless, the combination of IL4I1, SPP1, and NLRP3 antibodies was sufficient to detect and discriminate NLRP3 TAMs, SPP1 TAMs, and IL4I1 TAM group (encompassing ISG15 and CXCL9 TAMs that we could not resolve) that together labels all scRNA TAMs subsets (**Fig 3A**). Of note, the Proliferating TAMs are composed of a mixture of cells from different scRNA TAM clusters and form a separate cluster because their gene expression profiles are commonly enriched in cell cycle associated gene expression.

**Fig 3.**
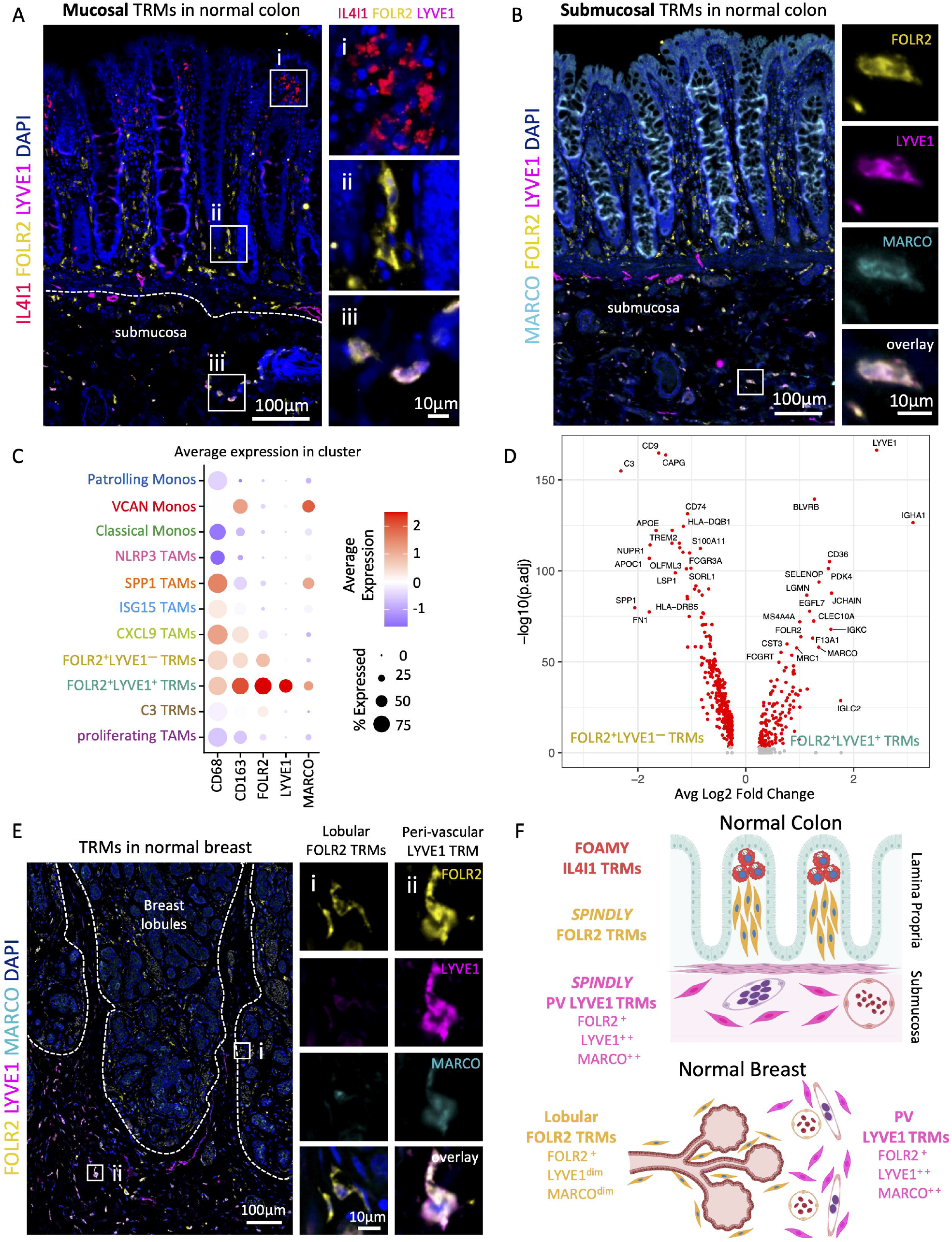
FOLR2, IL4I1, NLRP3, and SPP1 mark spatially distinct macrophage niches in the TME. (**A**) Dotplot shows average macrophage marker expression in scRNA macrophage populations and indicates which scRNA macrophage populations are detectable in 4-color IF staining by anti-NLRP3, -SPP1, -IL4I1, -FOLR2, and a combination of anti-FOLR2, -LYVE1 and -MARCO antibodies. (**B-E**) *Left:* CODEX image (**B**) or Immunofluorescence (IF) images (**C**,**D**,**E**) show the distribution of CD68 and CD163 (**B**), or FOLR2 and IL4I1 (**C**), NLRP3 (**D**), SPP1 (**E**) protein expression in representative cases of CRC (**B**,**C**,**E**) and BC (**D**). PanCK marks tumor cells. Close-up images on the bottom correspond to boxed regions on the top. *Top right:* Scatterplots show the distribution of CD68 Macs, CD163 Macs, FOLR2 TRMs, IL4I1 TAMs, NLRP3 TAMs, SPP1 TAMs corresponding to IF images on the left. *Bottom right:* Boxplots show the distance quantification of each macrophage to the closest tumor cell corresponding cells identified on IF images on the left. Pairwise comparisons were determined using a two-sided Wilcoxon rank-sum test on 1092 (**B**) 580 (**C**), 739 (**D**), and 203 (**E**) cells. (**F**) Distance (μm) of CD68 and CD63 macrophages to the closest tumor cell. Cells were identified on CODEX images. (**G**) Distance (μm) of IL4I1 TAMs, NLRP3 TAMs, SPP1 TAMs, FOLR2 TAMs to the closest tumor cell. *P values* were calculated with a linear mixed-effect model with Bonferroni’s corrections for multiple comparisons.

The four panels in **Fig 3B-E** show representative staining results of macrophage distribution in a single representative 1.5 mm^2^ tissue region of BC and CRC. Each panel shows 1) an IF image of the discussed markers (*left*), 2) a corresponding dotplot representing the spatial macrophage distribution revealed by the IF (*top right*), and 3) corresponding distance quantification to the closest tumor cell in that specimen (*bottom right*). We also show distance quantification across multiple regions and patient samples (**Fig 3F, G**). We started by analyzing the spatial distribution of the CD68 and CD163. The common view is that CD163-positive macrophages are of M2-type, and that they help tumor growth and metastasis (Rőszer 2015) and so are expected to localize close to the tumor. Surprisingly, contrary to the current view, we found that macrophages with higher CD163 expression (**Fig S3B**) localized further away from the tumor nests (**Fig 3B i, Fig 3F**; average distance of 74.5 μm) compared to macrophages with higher CD68 expression (**Fig S3B**) that infiltrated and tightly surrounded tumor nests (**Fig 3B ii, Fig 3F**; average distance of average 35.9 μm).

Next, we interrogated the spatial distribution of FOLR2, IL4I1, NLRP3, and SPP1. We found remarkable and unexpected segregation of these markers where FOLR2 expression was associated with benign tissue localized further away from the tumor. In contrast, IL4I1 (**Fig 3C**), NLRP3 (**Fig 3D**), and SPP1 (**Fig 3E**) expression were concentrated immediately adjacent to the tumor cells. To confirm this, we performed a distance comparison over 36,041 macrophages spanning 60 distinct 1.5 mm^2^ tissue fragments derived from 14 CRC and 13 BC cases. This analysis showed that IL4I1 TAMs were located an average of 38.3 μm away from the closest tumor cell, NLRP3 TAMs 47.4 μm away, SPP1 TAMs 36.4 μm away, and FOLR2 TRMs 109 μm away (**Fig 3G**).

### Spatial segregation of IL4I1 and FOLR2 macrophages is conserved across benign colon, primary tumor, and lymph node metastasis

Since we found remarkable spatial segregation of IL4I1 TAMs and FOLR2 TRMs in the benign colon (**Fig 2A**) and the TME in colon cancer (**Fig 3G-I**), we sought to investigate whether this pattern is conserved in the metastatic lesions. We compared IF staining of a benign colon, a CRC invasive front, and a lymph node CRC metastasis to show that IL4I1 TAMs and FOLR2 TRMs occupied spatially distinct histological regions (**Fig S3C-E**). We show that the distinct spatial segregation of IL4I1 from FOLR2 expressing macrophages observed in benign colon LP (**Fig S3C**) was conserved in the invasive front of CRC (**Fig S3D**) and LN CRC metastasis (**Fig S3E**). Similar to the invasive front of the CRC tumor, in the LN CRC metastasis, IL4I1 macrophages were present in the desmoplastic stroma surrounding the tumor nests, and FOLR2 macrophages were present further away in the surrounding benign tissue. In addition, we report that a thin buffer zone of macrophages expressing both FOLR2 and IL4I1 existed in both benign and tumor specimens, suggesting that local tissue cues, e.g. epithelial cell death, drive the IL4I1 and FOLR2 populations.

### IL4I1 marks phagocytosing macrophages in the benign colon, in the LN, and the TME

IL4I1 localizes in the lysosomes of antigen-presenting cells (Mason et al. 2004), suggesting a role in phagocytosis. A close inspection of the IF-stained benign colon mucosa revealed pan-cytokeratin (CK)-positive granules localized inside IL4I1 TRMs at the top of the colon crypt but not in FOLR2 TRMs in the middle and bottom of the crypt (**Fig 4A**). Our finding is consistent with work showing that macrophages ingest dying intestinal epithelial cells (IEC) at the top of the intestinal lamina propria (Nagashima et al. 1996), but provides a novel marker for this phenomenon. Next, we asked whether another specialized body phagocyte type, tingible body macrophages (TBMs), shows IL4I1 positivity. The TBMs localize in germinal centers where they remove apoptotic B cells (Aguzzi, Kranich, and Krautler 2014) and thus are expected to have a high expression of phagocytic markers. We found that they displayed very bright IL4I1 staining in both the LN germinal centers (**Fig 4B**) and the intestinal lymphoid follicles (data not shown). TBMs contain apoptotic cellular debris at different degradation stages and are named after apoptotic nuclear debris (‘tingible bodies’) that can be observed in their cytoplasm. A presence of TBMs rich in tingible bodies is a hallmark of follicular hyperplasia and Burkitt’s lymphoma, both characterized by fast cell turnover (Gotur and Wadhwan 2020). We examined hyperplastic LN from two different patients and two Burkitt’s lymphoma cases, and found that TBMs in both conditions contain many tingible bodies and display high IL4I1 expression (**Fig 4C-D**).

**Fig 4.**
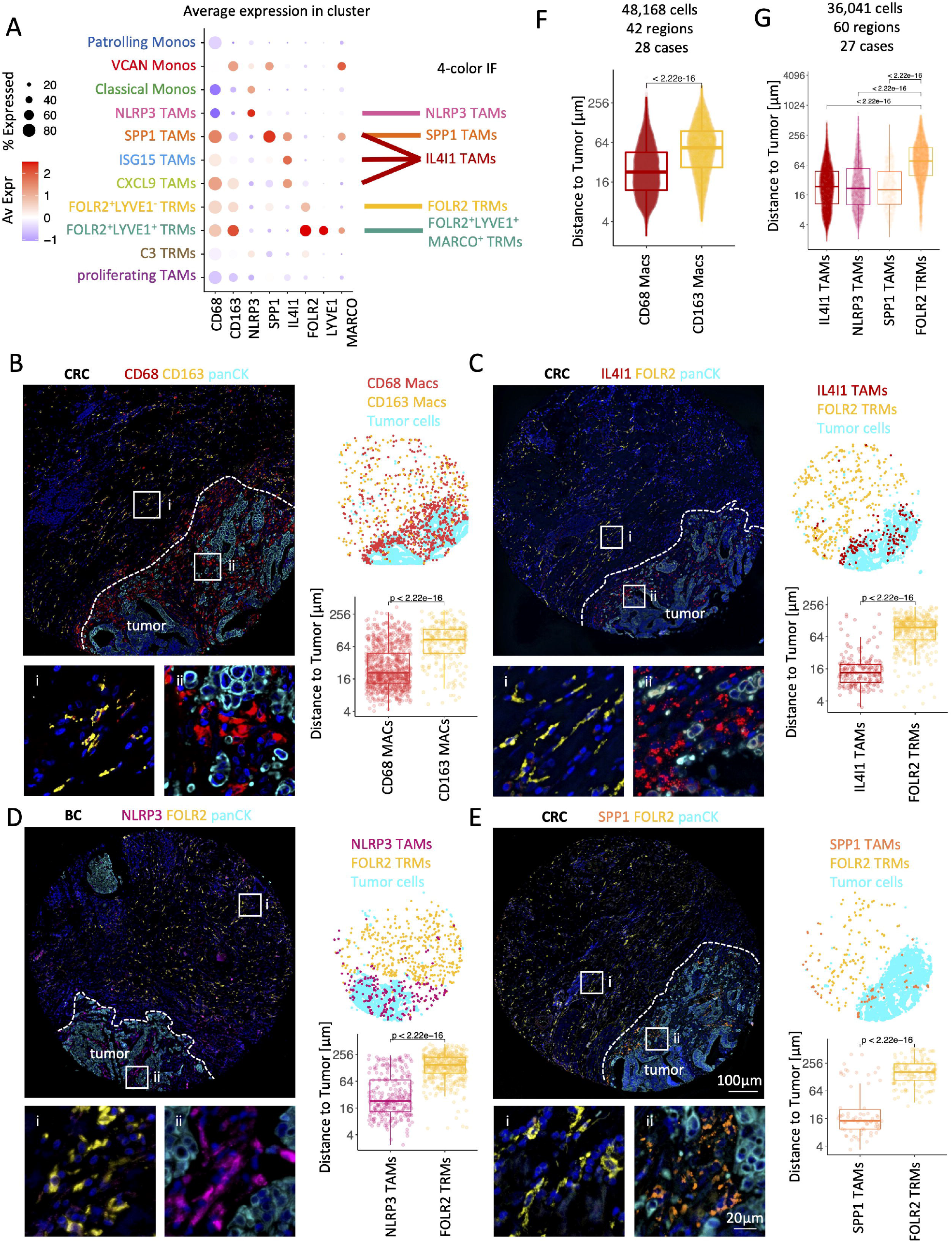
IL4I1 marks phagocytosing macrophages in the normal colon, in the LN and the TME. (**A**) IF images of normal colon mucosa stained with IL4I1, FOLR2, panCK, and DAPI show the presence of panCK^+^ material within IL4I1 macrophages. Close-up images on the right correspond to the boxed region on the left. (**B**) IF images of normal Lymph Node stained with IL4I1, FOLR2, and DAPI. (i) is a close-up image of a germinal center tingible body macrophage (TBM), (ii) is a close-up image of interfollicular FOLR2 TRMs. (**C**) Images of TBMs in hyperplastic lymph node and (**D**) Burkitt’s lymphoma staind with *top*: H&E and *bottom*: IL4I1 and DAPI. (**E**) Same as (**A**) but at the invasive front of CRC. (**F**) *Top:* KEGG pathways enrichment analysis of phagocytosis-related pathways across scRNA macrophage populations. Populations with no significantly enriched pathways were omitted. *Bottom:* average IL4I1 gene expression across scRNA macrophage populations with enriched phagocytosis-related gene sets. (**G**) Dotplot shows average gene expression in scRNA macrophage populations. (**H**) Barplots show frequency of scRNA monocyte and macrphage clusters from Bassez et al., dataset stratified by response to pembrolizumab and time of sample collection. (**I**) Boxplots show frequency of scRNA monocyte and macrophage clusters pre pembrolizumab treatment from Bassez et al. (**J**) Schematic illustrating IL4I1 TAM association with cell death and efferocytosis and highlighting IL4I1 TAMs as potential anti-CD47 (indirect as IL4I1 TAMs express CD47 ligand-SIRP1?) and anti-PD-L1 (direct) therapy targets.

Given these results, we next asked whether we could detect evidence of tumor cell phagocytosis in the TME, and found that IL4I1 TAMs in the invasive front of the tumor also contained pan-CK granules we interpret as apoptotic bodies derived from tumor cells (**Fig 4E**). Subsequently, we used gene set enrichment analysis to inquire whether scRNA IL4I1 positive scRNA macrophage subsets are enriched in phagocytic processes. We found that compared to other scRNA macrophage subtypes, the CXCL9 TAMs (a subset of *IL4I1*^*+*^ TAMs) were most enriched in Phagosome, Lysosome, Endocytosis, and Antigen Processing and Presentation gene sets expression (**Fig 4F**).

Next, we asked whether IL4I1 TAMs might be targets of phagocytosis-modulating cancer therapies including anti-CD47 and anti-PD-L1 treatment (Gordon et al. 2017). Notably, using our integrated myeloid object (**Fig 1C**) we found that *SIRPA* encoding SIRPα-a CD47 ligand, and *CD274* encoding PD-L1 were both enriched in *IL4I1* expressing scRNA myeloid clusters including SPP1 TAMs, ISG15 TAMs and CXCL9 TAMs (**Fig 4G**). This indicates that IL4I1 TAMs are indirect anti-CD47 and direct anti-PD-L1 immunotherapy targets. Thus next, we asked whether IL4I1 TAMs could be used as predictive marker of PD-1–PD-L1 axis blockade. For this analysis we used the scRNA monocyte and macrophage transcriptomes form Bassez et al., dataset (**Fig 1C, S1A-B**) that contains samples of advanced breast cancer patients taken before and after pembrolizumab treatment. We found that the frequency of IL4I1 expressing scRNA TAMs, both pre- and post-treatment, was increased in patients that responded to the therapy (**Fig 4H-I**).

These results demonstrated that IL4I1 is a marker associated with active phagocytosis in benign tissue and malignancy and suggest that IL4I1 TAMs might be implicated in anti-CD47 and anti-PD-L1 immunotherapy response (**Fig 4J**), and serve as a predictive marker of PD1–PD-L1 axis blockade.

### CODEX multiplexed imaging reveals cellular co-enrichment in macrophage niches

Having identified the spatial segregation of the IL4I1, NLRP3, SPP1, FOLR2, and LYVE1 macrophage populations, we next sought to elucidate the cellular compositions of the spatially segregated macrophage niches. We used CO-Detection by indEXing (CODEX) multiplexed tissue imaging to simultaneously visualize 36 protein markers on a single tissue microarray section of breast and colon benign and tumor tissue (Black et al. 2021; Kennedy-Darling et al. 2021; Goltsev et al. 2018). Our CODEX antibody panel contained four canonical myeloid markers (CD16, CD68, CD163, CD206). To further subtype the macrophage populations, we added SPP1, LYVE1, and FOLR2. Using the CODEX computational pipeline (i.e., imaging processing, single-cell segmentation, and unsupervised clustering) (Hickey, Tan, et al. 2021), we identified two epithelial cell types, seven stromal cell types, and fifteen immune cell types (**Fig 5A, Fig S4A**). Among the immune cell types, we discriminated five macrophage subsets: CD68 TAMs, SPP1 TAMs, CD163 TRMs, FOLR2 TRMs, and LYVE1 TRMs (**Fig S4B**). The CODEX-identified CD68 TAMs likely corresponded to IL4I1 TAMs and NLRP3 TAMs populations we identified by IL4I1 and NLRP3 staining in our IF studies. For technical reasons, we were unable to add IL4I1, NLRP3, and MARCO antibodies to the CODEX panel. Thus, the CODEX-identified CD163 TRMs might represent LYVE1 TRMs and FOLR2 TRMs.

**Fig 5.**
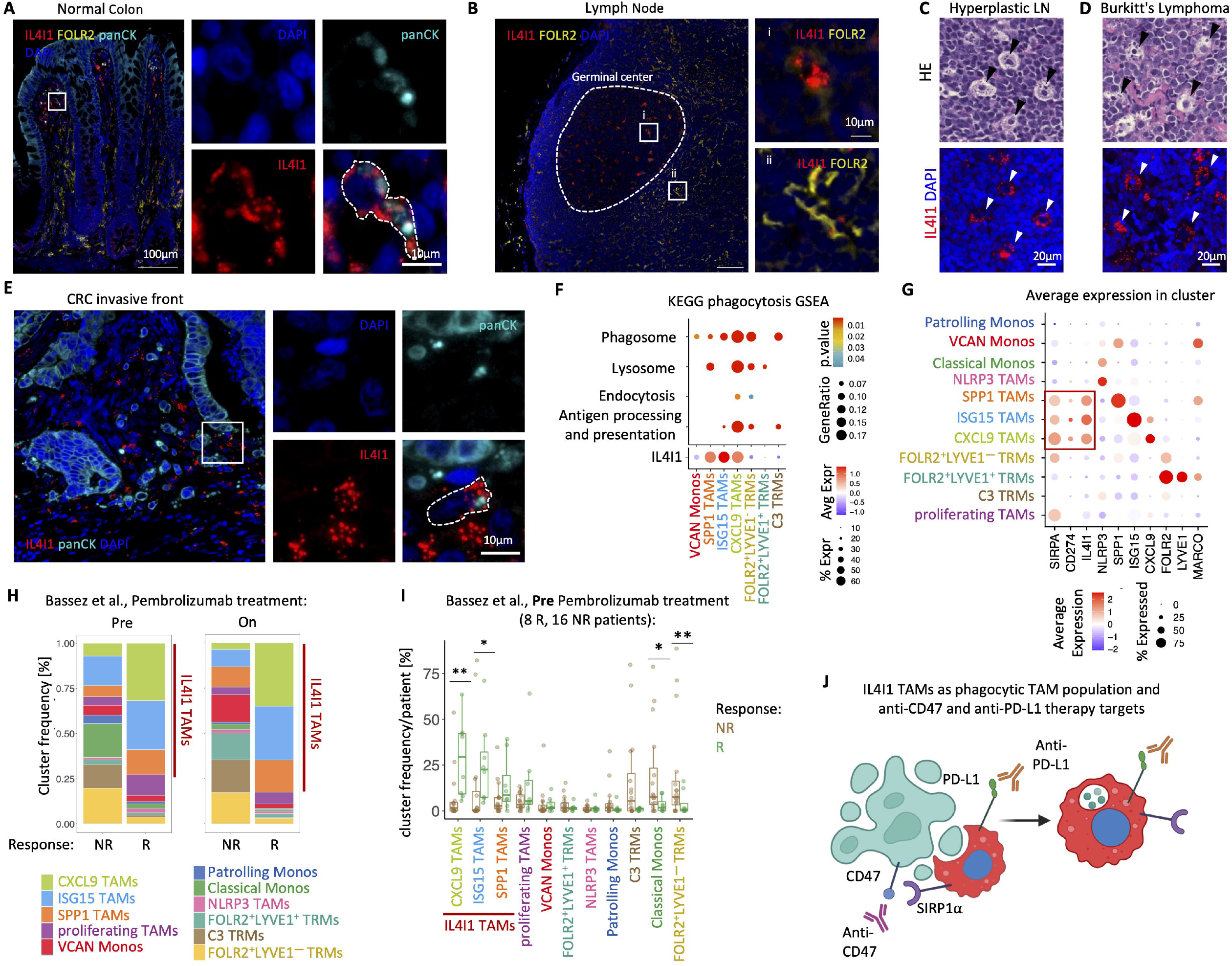
36-antibody CODEX multiplexed imaging reveals cellular interactions in macrophage niches within colon and breast cancer tissues. (**A**) Schematic shows CODEX imaging and cellular neighborhood analysis workflow. (**B**) Heatmap shows CODEX cell types (x axis) enrichment (color) in the identified cellular niches (y axis). (**C**) Boxplot shows distance (μm) to the closest tumor cell for every macrophage identified by CODEX labeled by the niche it belongs to. (**D**) Barplot shows a percentage of the epithelial cells occupied in each CODEX macrophage niche. (**E**) Barplot presents the frequency of CODEX macrophage niches grouped by anatomical location. NB-normal breast, DCIS-ductal carcinoma in situ breast, IDC-invasive ductal carcinoma breast, NGI-normal GI tract, IF invasive front CRC, CT-center of tumor CRC. (**F**) Schematic shows cellular macrophage niche organization and closeness to the tumor.

CODEX imaging showed that the distribution of CD68 and CD163 is different between the five macrophage subsets, with CD68 and SPP1 TAMs enriched in CD68 expression while CD163, FOLR2, and LYVE1 TRMs enriched in CD163 expression (**Fig S4B**). Consistent with the scRNA Seq and 4-color IF results (**Fig 1D, Fig 3B**), CODEX imaging confirmed the existence of 2 FOLR2 positive macrophage populations: FOLR2^+^LYVE1^−^ and FOLR2^+^LYVE1^+^ (**Fig S4B**). Moreover, we validated that SPP1 TAMs (average distance 28.4 μm) localize closer to the tumor than FOLR2 macrophages (average distance 65.8 μm) (**Fig S4C**). In addition, CODEX data showed that similar to FOLR2 TRMs, the CODEX LYVE1 TRMs are localized further away from the tumor (average distance 106 μm) (**Fig S4C**).

To uncover the cellular composition of the spatial macrophage niches, we next performed cellular neighborhood analysis on the CODEX multiplexed imaging data (Schürch et al. 2020; Phillips et al. 2021; Jiang et al. 2022). We clustered cells based on the identity of their ten closest neighboring cells and identified 14 cellular niches, of which nine were enriched in macrophages (**Fig 5A**). We grouped the nine macrophage-containing niches into four niche types, each named after the primary macrophage subtype it contains: 1) CD68 TAM niche, 2) SPP1 TAM niches, 3) FOLR2 TRM niches, and 4) LYVE1 TRM niche (**Fig 5B**). The one CD68 TAM niche was localized inside the tumor nests and co-enriched with the tumor cells (**Fig 5B, S5A**); we called it the *Intra-tumoral TAM niche*. The three discrete SPP1 TAMs niches were all enriched with SPP1 TAMs and the tumor cells but differed in cellular composition. The *Peri-tumoral SPP1 TAM Niche* contained CD68 macrophages (**Fig 5B, S5B**), the *Inflamed SPP1 TAM Niche* contained neutrophils (**Fig 5B, S5C**), and the *Hypoxic SPP1 TAM Niche* held hypoxic tumor cells marked by CA9 expression (**Fig 5B, S5D**). The four discrete FOLR2 niches were co-enriched in FOLR2 TRMs and CD163 TRMs but had different cell compositions and tissue locations. *The Plasma Cell (PC) enriched FOLR2 TRM Niche* was co-enriched with PCs and located close to the blood vessels and in the normal gastrointestinal (NGI) LP (**Fig 5B, S5E**). *The Smooth Muscle FOLR2 TRM Niche* labeled the bowel muscle wall (**Fig 5B, S5F**). *The Trapped Fibrous FOLR2 TRM Niche* was enriched in FAP fibroblasts and marked fibrous bands entrapped between growing tumor nests (**Fig 5B, S5G**). The *Lymphoid FOLR2 TRM Niche* was composed of CD4T, CD8T, Tregs, DCs, and FOLR2 TRMs (**Fig 5B, S5H**). The LYVE1 TRM niche was co-enriched with LYVE1 TRMs, FOLR2-TRMs, CD163 TAMs, PDGFRb fibroblasts, mast cells, and blood and lymph vessels. We called it the *Peri-Vascular LYVE1 TRM Niche* (**Fig 5B, S5I**).

Next, we used two approaches to map each CODEX-macrophage niche tissue distribution relative to the tumor. First, we computed the distance of every macrophage, labeled by the niche it belongs to, to the closest tumor cell (**Fig 5C**). Second, we calculated the fraction of tumor cells in every macrophage-enriched niche (**Fig 5D**). We interpret the distance to the tumor and the fractional enrichment in tumor cells as an indicator of how closely the given niche is associated with the tumor. These analyses revealed a remarkable spatial macrophage niche segregation and a 3-tier distribution of closeness to the tumor. Specifically, we show that TAMs in *the Hypoxic SPP1 Niche* and *the Intra-tumoral Niche* were located the closest to the tumor cells with an average distance of 9.37 and 10.6 μm to the nearest tumor cell (**Fig 5C**) and that those two niches had the highest fraction of tumor cells (**Fig 5D**). In contrast, TRMs in *the Lymphoid FOLR2, the PCs enriched FOLR2, the Peri-Vascular LYVE1* and *the Smooth Muscle FOLR2 Niches* lay the farthest from the tumor with an average distance of 55.2, 57.5, 74.8, and 76.0 μm from the closest tumor cell (**Fig 5C**). In agreement, they also contained the smallest percentage of tumor cells (**Fig 5D**). Macrophages in *The Peri-tumoral SPP1, the Inflamed SPP1*, and *the Trapped Fibrous FOLR2 Niches* localized at an intermediate distance between the two extremes.

To better visualize the spatial distribution of the macrophage niches in benign and tumor tissues, we plotted the niche frequency by anatomic location. We show that *the Peri-Vascular LYVE1 TRMs Niche* was most enriched in benign breast, while *the PCs enriched FOLR2 TRMs Niche* was most enriched in NGI mucosa. *The Smooth Muscle FOLR2 TRMs Niche* labels bowel wall and was thus specific to gut samples, and it could be detected in benign, in the invasive front and center of the tumor samples. This is consistent with the fact that CRC invades the bowel wall. In turn, the *Intra-tumoral TAM Niche, the Inflamed SPP1 TAM Niche, the Peri-tumoral SPP1 TAM Niche, the Hypoxic SPP1 TAM Niche, and the Trapped Fibrous FOLR2 Niche* were enriched in ductal carcinoma in situ (DCIS), invasive ductal carcinoma (IDC), in the IF of CRC and the CRC center of the tumor (CT), further supporting that they are tumor-associated (**Fig 5E**).

Taken together, the CODEX data (**Fig 5, S4, S5**) allowed us to identify spatial associations between macrophage subtypes and other cell types in benign and tumor tissues. We showed that SPP1 TAMs were co-enriched with CD68 TAMs close to the tumor cells, localized in hypoxic tumor areas, and associated with neutrophilic infiltration. In contrast, CD163 TRMs, FOLR2 TRMs, and LYVE1 TRMs were co-enriched in adjacent benign tissue located further away from the tumor. We showed that FOLR2 TAMs constituted a tissue-resident macrophage population in the bowel muscle wall and were associated with PCs in the intestinal lamina propria and connective breast tissue. We found that FOLR2 TRMs from the breast connective tissue or muscle bowel wall can be trapped within growing tumor nests and thus become a part of the TME (**Fig 5F**).

### FOLR2 macrophages and plasma cells spatially colocalize

To further explore the CODEX-identified FOLR2 TRM association with PCs, we used IHC and multicolor IF. Single color IHC showed that FOLR2 TRMs were in direct contact with PCs, histologically identified by the clock-like nuclear chromatin condensation pattern and asymmetric cytoplasmic ‘hof’ where antibodies are produced and stored (arrowheads, **Fig 6A**). Multicolor IF showed that FOLR2 TRMs and CD38^+^ PC occupied the same space in the middle and bottom layers of the colon lamina propria (**Fig 6B top panel**), corroborating the CODEX results. Furthermore, we found Lobular FOLR2 TRMs were immediately adjacent to PCs around benign breast glands (**Fig 6B bottom panel**). Previous studies showed that CD163^+^ macrophages surround PCs in the extrafollicular foci in the tonsil (Xu et al. 2012). Here we showed that it was the FOLR2 TRM subtype that localized directly next to PCs in the interfollicular zone in the LN (**Fig S6A**). Apart from PCs, CD38 can also be expressed by activated T cells, Monocytes, NK cells, and other immune cells (van de Donk et al. 2016). To unequivocally demonstrate that the CD38^+^ cells spatially co-enriched with FOLR2 TRMs were PCs, we used 4-plex IF staining and showed that cells localized directly next to FOLR2 TRMs were marked by overlapping expression of CD38 and a prototypical PC marker - CD138 (**Fig S6B**).

**Fig 6.**
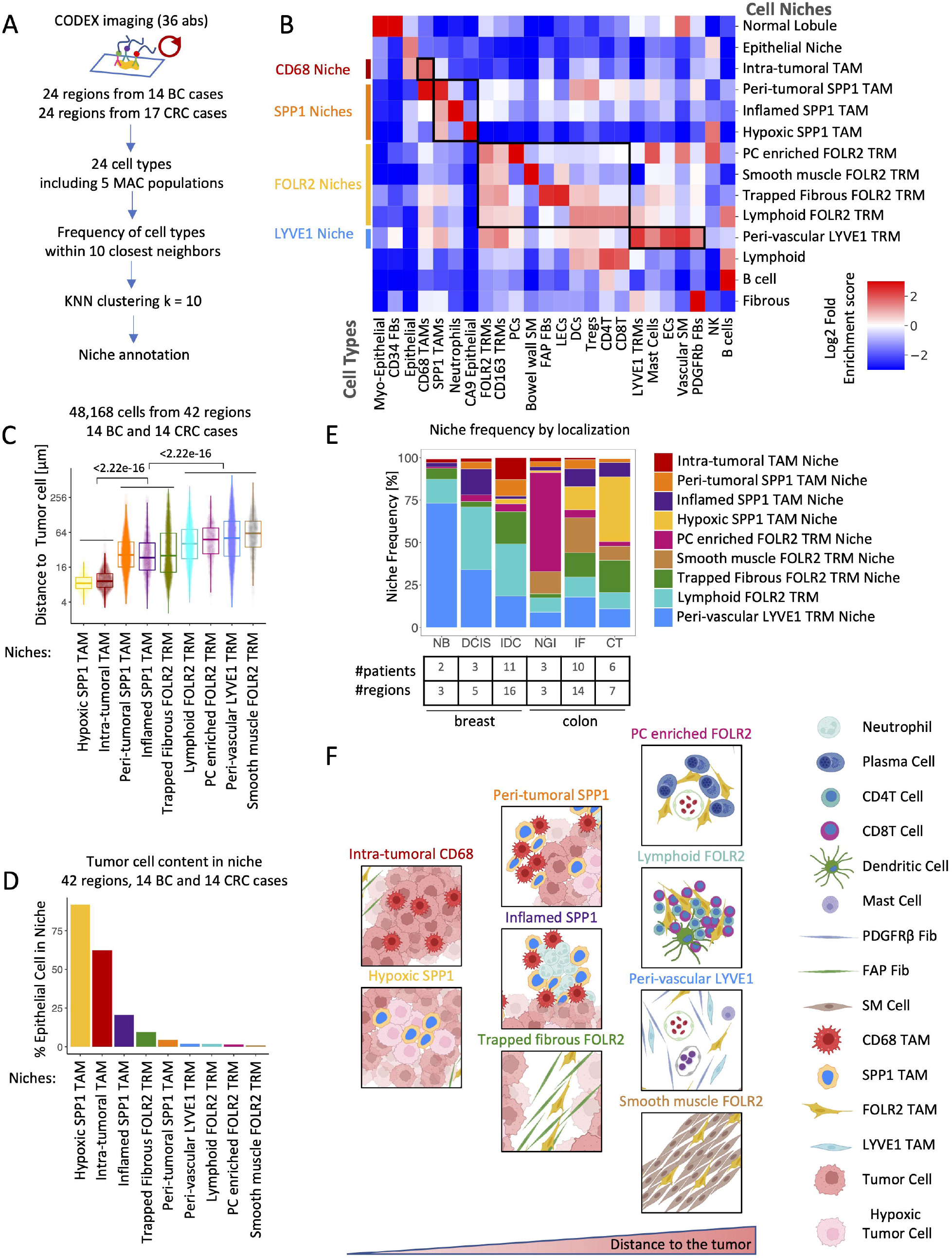
FOLR2 Macrophages and Plasma Cells spatially colocalize. (**A**) Immunohistochemical image shows FOLR2 TRMs surrounded by plasma cells indicated with black arrows. (**B**) Immunofluorescence (IF) images show FOLR2 TRMs colocalization with CD38 PCs on *Top:* the middle and bottom of the colon lamina propria and *Bottom:* normal breast gland. Middle: Close-up images in the middle correspond to boxed regions on the top and bottom IF images. The scale bar of 20 μm is identical for both close-up images. (**C**) Scatterplots show the distribution of FOLR2 TRMs, IL4I1 TAMs, and PC identified by CD138 staining in BC TME. (**A**-**B**) The scale bar of 10 μm is identical for all close-up images. (**D**) Boxplot shows distance quantification of each FOLR2 TRMs, IL4I1 TAMs to the closest tumor cell measured across 7 1.5 mm^2^ tissue regions of BC and CRC. *P* value calculated with a linear mixed-effect model. (**E**) Dotplot shows communication probability between all significant Ligand and Receptor interactions between FOLR2 TRMs and PCs in BC scRNA Seq dataset of Basses et. al. (**F**) Schematic illustrating possible FOLR2 TRMs interaction with PCs.

Next, to demonstrate that the association between PCs and FOLR2 TRMs was specific, we computed the distance from every IL4I1 TAM and FOLR2 TRM to their closest PC across seven different tissue regions. As anticipated, PCs clustered closer to FOLR2 TRMs than the IL4I1 TAMs (**Fig 6C-D**).

To interrogate the interaction mechanism between PCs and FOLR2 TRMs, we performed a scRNA Seq-based ligand-receptor interaction analysis. First, we used PCs and FOLR2 TRMs transcriptomes from the scRNA Seq study on BC patients (Bassez et al. 2021). The highest probability interactions were found between APRIL (TNFSF13) and BAFF (TNFSF13B) on the FOLR2 TRMs and BCMA (TNFRSF17) on the PCs (**Fig 6E**). APRIL and BAFF are known to drive PC infiltration and long-term survival in the tissue (Kawakami et al. 2019; Benson et al. 2008). Similarly, using the IgA^+^PC, IgG^+^PC, and FOLR2 TRMs scRNA Seq transcriptomes from benign colon and CRC (Lee et al. 2020), we identified BAFF (TNFSF13B) and BCMA (TNFRSF17) interaction as the highest probability interaction between IgA^+^PC and FOLR2 TRMs (**Fig S6C**). Our results are in agreement with previous literature that suggests antigen-presenting cells maintain a PC niche in human tonsils (Xu et al. 2012), murine bone marrow (Rozanski et al. 2011), and human lamina propria (Hickey, Becker, et al. 2021). Taken together, these observations suggest that FOLR2 TRMs play a key role in recruiting and maintaining PCs in benign colon lamina propria and inflamed tumor-adjacent benign tissue (**Fig 6F**).

### NLRP3 and SPP1 TAMs are associated with neutrophils in the TME

CODEX neighborhood analysis revealed spatial co-enrichment of SPP1 TAMs with neutrophils in *the Inflamed SPP1 TAM niche*. Notably, we also found NLRP3 TAMs to be enriched in neutrophil infiltrated areas (**Fig 7A**). However, unlike NLRP3 TAMs, which were spatially co-enriched with live neutrophils, SPP1 TAMs were associated with necrotic tissue areas (**Fig 7B**). This observation prompted us to compare NLRP3 and SPP1 TAMs’ transcriptomes. Differential gene expression showed that NLRP3 TAMs expressed high levels of neutrophil chemoattractant cytokines (*CXCL1, CXCL2, CXCL8*). In contrast, the most upregulated genes in SPP1 TAMs were associated with phagocytosis and lipid metabolism, including apolipoproteins (*APOC1, APOE*), lipid scavenger receptors (*TREM2, MARCO*), lipid transporter *FABP5*, cathepsins (*CTSB, CTSD, CTSZ*), and matrix metalloproteinase (*MMP9, MMP12*) (**Fig 7C-D**). Interestingly, SPP1 itself has been implicated in phagocytosis (Shin et al. 2011; Schack et al. 2009) and lipid metabolism (Remmerie et al. 2020). To further interrogate the association of SPP1 TAMs with necrosis, we used a publicly available 10x Visium FFPE Human Breast Cancer sample to show that necrotic tumor areas were enriched in SPP1 rather than NLRP3 gene expression (**Fig S7A-B**). These results suggest that NLRP3 TAMs likely contribute to neutrophil recruitment in the TME and that the SPP1 TAMs may play a role in the phagocytosis of the dying cell in the necrotic tumor areas.

**Fig 7.**
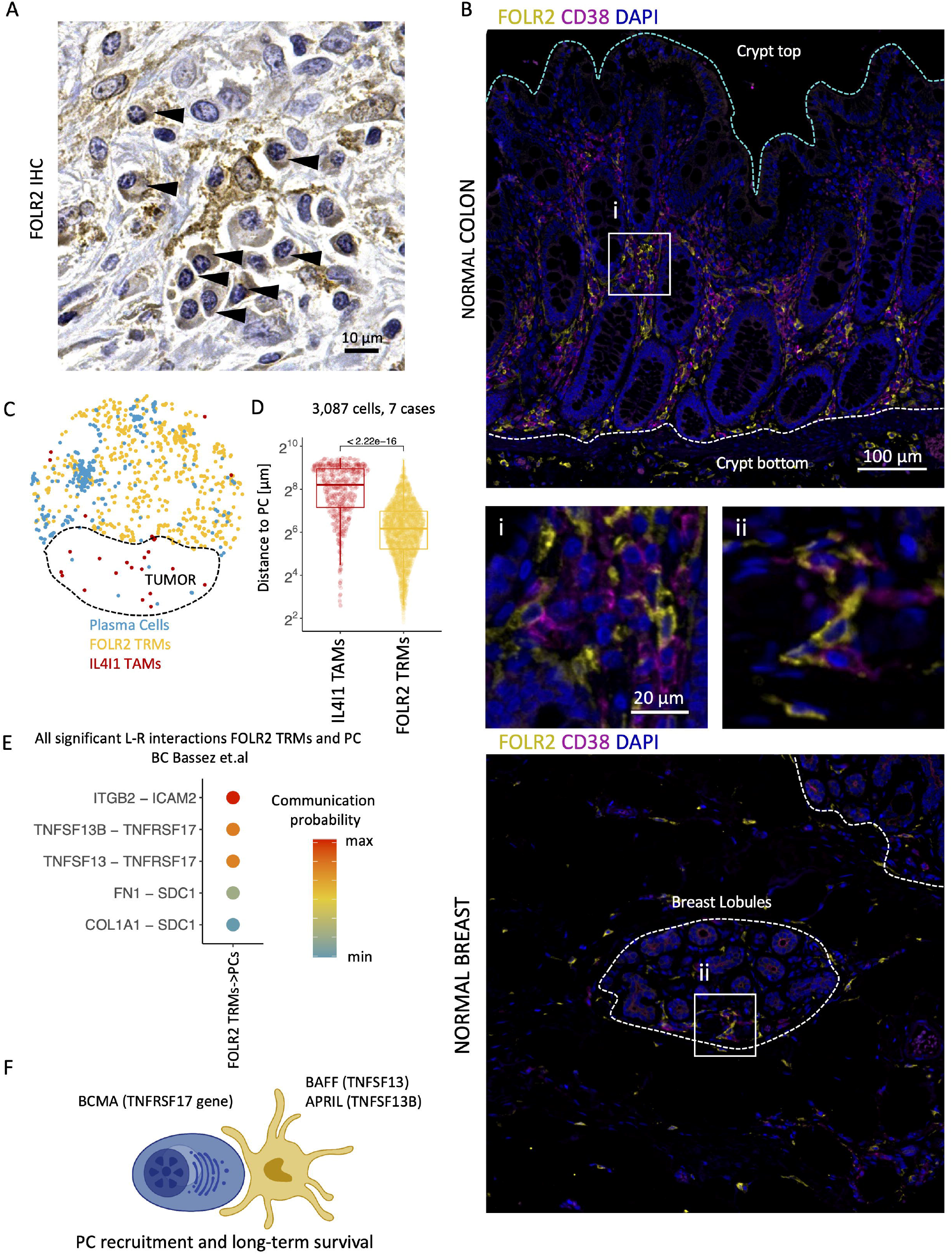
NLRP3 and SPP1 TAMs are associated with neutrophils in the TME. **(A)** Immunohistochemical image shows NLRP3 TRMs surrounded by neutrophils (arrowheads). **(B)** Immunohistochemical image shows SPP1 TRMs surrounded by karyorrhectic debris in necrotic material (arrowheads). (**C**) Volcano plot shows differential gene expression between scRNA transcriptomes of SPP1 TAMs and NLRP3 TAMs. (**D**) Dotplot of average expression of genes associated with neutrophil chemoattraction, lipid metabolism and phagocytosis across scRNA macrophage populations. (**E**) Immunofluorescence (IF) shows a representative BC regions stained with NLRP3, CD68, Calprotectin (CPTN) and DAPI. Scale bar of 10 μm is identical for all close-up images. (**F**) Quantification of the number of neutrophils present on 9 BC 1.5 mm^2^ tissue regions stratified by whether they contained diffuse NLRP3 (3 regions) or NLRP3 specks (6 regions). *P* value was computed using a two-sided Wilcoxon’s rank-sum test. (**G**) IF images show CD68, CPTN and DAPI staining of a representative region of macrophage infiltrate in Crohn’s Disease. *Bottom:* Close-up images on the bottom correspond to boxed regions on the top IF images and show NLRP3, CD68, CPTN and DAPI staining. (**H**) Quantification of the number of neutrophils present in 9 normal colon submucosa and 9 macrophage infiltrated Crohn’s Disease areas with NLRP3 specks. *P* value was computed using a two-sided Wilcoxon’s rank-sum test. (**I**) Schematic of a possible mechanism through which NLRP3 TAMs can contribute to the recruitment of neutrophils in the TME. (**J**) Survival associations of single gene or macrophage niche signatures stratified by tumor type.

NLRP3 is a pathogen and danger-associated molecular pattern receptor known to form an intracellular complex called the inflammasome that leads to proteolytic IL1β activation and release. IL1β is known to play a role in neutrophil recruitment in infection (Miller et al. 2007) and cancer (Chen et al. 2012). Interestingly, we observed that the NLRP3 expression could be either diffuse (**Fig 7E i**) or aggregated in a speck, a micrometer-sized structure formed by the inflammasome (**Fig 7E ii**). We found that speck-like NLRP3 aggregation, which we interpret as activated inflammasome complexes, was linked to neutrophil infiltration (**Fig 7E ii**). To confirm, we stratified BC and CRC positive regions by whether they were enriched in NLRP3 macrophages with diffuse staining or contained TAMs with NLRP3 specks, and quantified the number of neutrophils; NLRP3 specks correlated with neutrophil presence (**Fig 7F**). Thus, we hypothesize that NLRP3 inflammasome assembly in the NLRP3 TAMs induces IL1β activation and secretion, which drives neutrophil infiltration (**Fig 7I**).

To extend our findings beyond cancer, we investigated whether we could detect NLRP3 inflammasome activation in Crohn’s Disease (CD), a type of inflammatory bowel disease associated with neutrophil infiltration. Indeed, the examination of three cases of advanced CD showed that regions with high macrophage infiltration 1) contained NLRP3 specks and 2) were highly infiltrated by neutrophils (**Fig 7G**). Comparing neutrophil quantification from nine regions in three CD cases to nine regions in the benign submucosal colon showed that neutrophil infiltration was associated with NLRP3 inflammasome aggregation in CD (**Fig 7H**). While previous studies suggested inflammasome involvement in cancer (Ridker et al. 2017; Sharma and Kanneganti 2021) and CD (Villani et al. 2009), we were the first to visualize the actual inflammasome formation in human BC, CRC, and CD in human FFPE tissue sections and to demonstrate an association of the NLRP3 inflammasome formation with neutrophil infiltration.

Previous reports showed that macrophage subtype signatures, including SPP1 TAM (Zhang et al. 2020; Lee et al. 2020) and FOLR2 TRM (Ramos et al. 2022), predict clinical cancer outcomes. However, the association of macrophage niches with clinical outcomes remains largely unexplored. We, therefore, determined the prognostic association of NLRP3 TAM and SPP1 TAM neutrophil niches in clinically-annotated datasets, including the PRECOG data (Gentles et al. 2015) (STAR Methods). In addition, we interrogated single marker gene associations, and a FOLR2/SEPP1/SLC40A1 gene signature, previously associated with favorable clinical outcomes in BC, as a reference. This analysis showed that both *SPP1* gene expression and abundance of *the Neutrophil inflamed SPP1 TAM Niche* were very strong predictors of poor prognosis in BC and CRC. Surprisingly, we found that *NLRP3* gene expression alone did not correlate with BC or CRC patient outcomes, while the abundance of *the Neutrophil inflamed NLRP3 TAM Niche* was strongly associated with adverse BC outcomes. Moreover, in line with previous reports (Ramos et al. 2022), we found that the FOLR2/SEPP1/SLC40A1 signature predicted favorable outcomes in BC but not CRC (**Fig 7J, S7C-D**).

Taken together, these results suggest that NLRP3 TAMs may be involved in the onset of inflammation through activating the NLRP3 inflammasome and may be driving neutrophil infiltration in TME and Crohn’s Disease. In addition, we demonstrate that the abundance of NLRP3 TAMs and neutrophil niche is associated with poor BC patient outcomes, suggesting that NLRP3 targeting in cancer might be a novel and promising treatment avenue.

## Discussion

This work reveals a rich landscape of spatially segregated functional macrophage niches across healthy and malignant human breast and colon tissue. We demonstrate that macrophage niches are not specific to an anatomical location or disease but rather conserved between tissue compartments with similar local cues. For example, IL4I1 macrophages are embedded in areas enriched in cell death like the colonic upper lamina propria, LN germinal centers, and desmoplastic stroma at the invasive front of the tumor. Thus, our findings indicate that macrophage niches are fundamental functional building blocks of tissue. In addition, we uncover some of the incoming and outgoing signals governing the macrophage niche. For example, we demonstrate NLRP3 inflammasome activation in *the Neutrophil inflamed NLRP3 TAMs Niche* and propose that it might result in IL-1β release and subsequent neutrophil recruitment.

It has been recognized that TRMs across different organs exhibit specialized functions reflecting local tissue physiology (Okabe and Medzhitov 2016). However, we are the first to uncover the existence of distinct functional spatial niches harboring discrete macrophage populations and cellular compositions within a single organ system. In particular, we reveal the existence of four separate macrophage niches in the bowel wall, including a phagocytic IL4I1 TAM niche, a novel FOLR2 TRMs plasma cell niche, a peri-vascular LYVE1^+^FOLR2^+^ TRMs niche in the bowel submucosa, and a smooth muscle FOLR2 TRMs niche in the muscularis propria.

Notably, our results reveal that IL4I1, SPP1, and NLRP3 TAM niches are closely associated with the tumor nests and implicated in the cancer response, including tumor cell death, hypoxia and tissue necrosis, and acute inflammation, respectively. In addition, IL4I1 TAMs might be implicated in response to anti-CD47 and PD1-PD-L1 blocking therapy as they express CD47 ligand-*SIRPA* and *CD274* encoding PD-L1, and correlate with anti-PD1 treatment response. Moreover, we show that NLRP3 inflammasome activation correlates with acute inflammation in BC, CRC, and CD and is associated with adverse patient outcomes in BC. This finding nominates the NLRP3 inflammasome as a novel therapy target and its specific small molecule inhibitor - MCC950 (Coll et al. 2015), as a novel therapeutic agent in solid tumors and CD.

Collectively, our findings elucidate a landscape of discrete human macrophage niches, uncover unexpected cell interactions and mechanisms governing the macrophage niche biology, explore the prognostic significance, and suggest novel therapy targets. Importantly, since the tools we present are FFPE-compatible, they enable the use of archival clinical material and provide a framework for the study of human macrophage function in health and disease.

## Limitations of the study

Ideally, macrophage tissue distribution and function should be profiled by simultaneous visualization of all macrophage populations. However, we could not include IL4I1 and NLRP3 antibodies for CODEX imaging due to incompatibility of working FFPE clones with DNA tags for adequate staining. In effect, we detected a large population of CODEX CD68 TAMs that localize close to the tumor and likely correspond to the IL4I1 and NLRP3 TAMs we characterize using the IF. Additionally, the evidence we present to propose the function of the discrete macrophage population is based on gene expression and imaging observations. Thus our findings warrant and inform future functional studies that should validate our observations.

## Figure Legends

**Fig S1.**
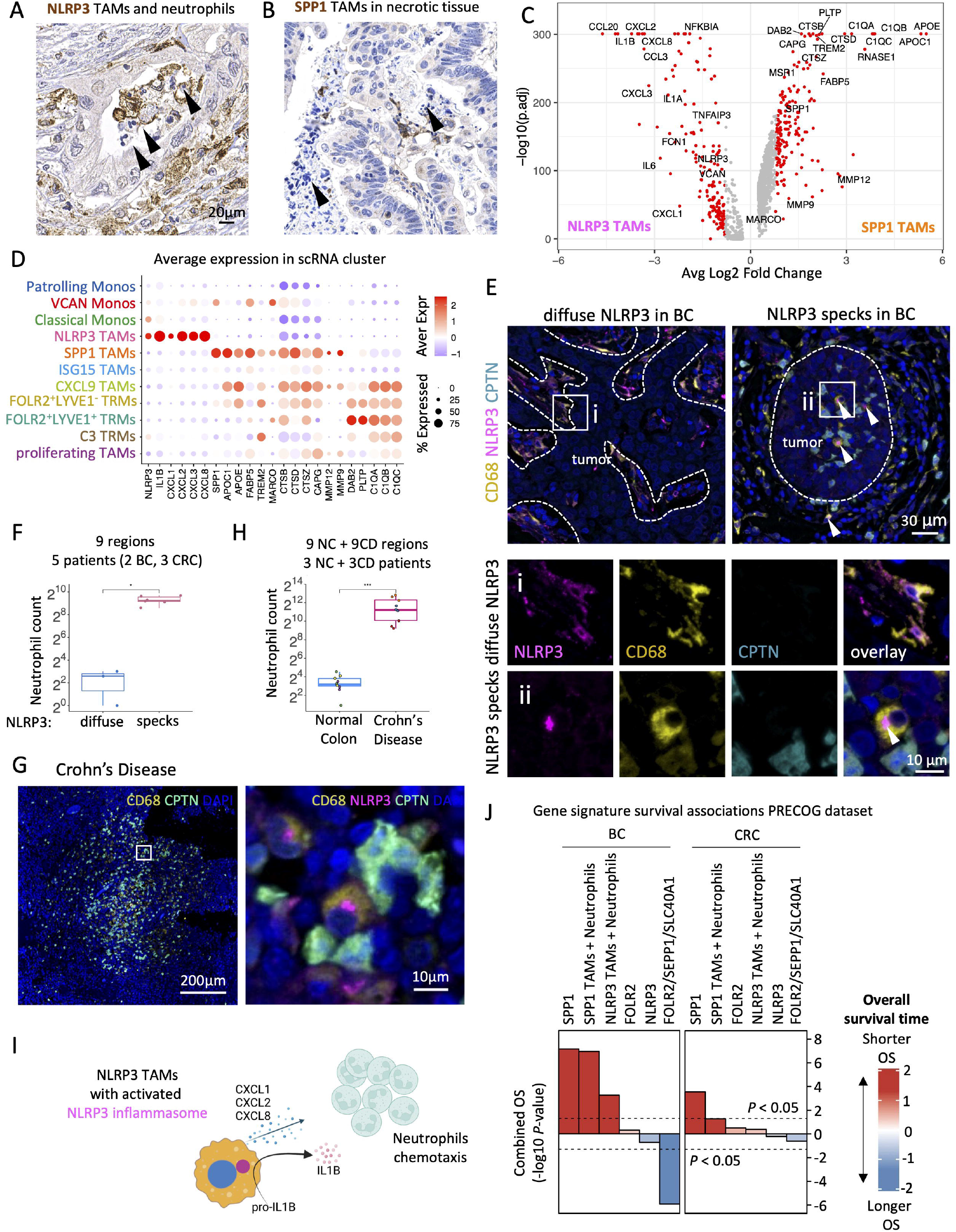
Single-cell RNA Sequencing and spatially resolved transcriptomics reveal differences in spatial enrichment of myeloid markers, related to Fig 1. (**A**) UMAP projection of monocytes and macrophages scRNA transcriptomes grouped by and colored by dataset showing the contribution of each dataset. (**B**) UMAP projection of monocytes and macrophages scRNA transcriptomes colored by tumor type. (**C**) Boxplots show the frequency of scRNA macrophage populations across 37 samples in 31 CRC patients ordered by their average expression. (**D**) Same as (**C**) but in 48 BC patients. (**E**) Same as (**C**) but in 8 normal colon samples and 37 CRC samples in 31 CRC patients and ordered by average frequency of cell populations in Tumor samples.

**Fig S2.**
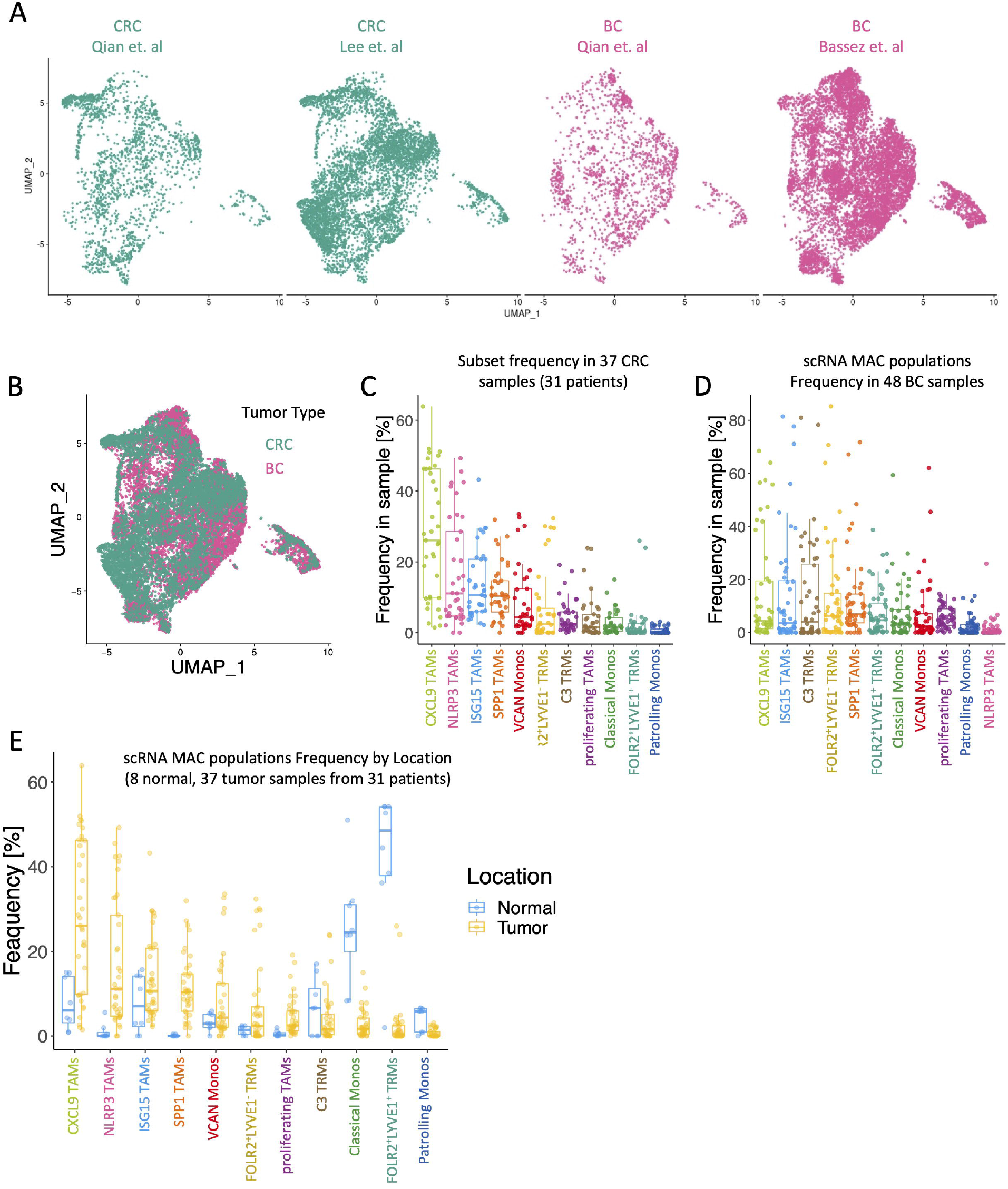
IL4I1, FOLR2, LYVE1 label spatially segregated TRM niches in human gastrointestinal tract, related to Fig 2. (**A**) Immunofluorescence (IF) images show CD68 and CD163 distribution in normal colon mucosa (i), colon lymphoid follicle (ii), and submucosa (iii). (**B**) IF image shows 3 TRM layers marked by IL4I1 (i), FOLR2 (ii), and FOLR2 and LYVE1 (iii) in normal duodenum mucosa and submucosa. (**C**) same as C but in normal Ileum mucosa and submucosa. (**A-C**) Close-up images on the right correspond to boxed regions on the left. The scale bar of 10 μm is identical for all close-up images.

**Fig S3.**
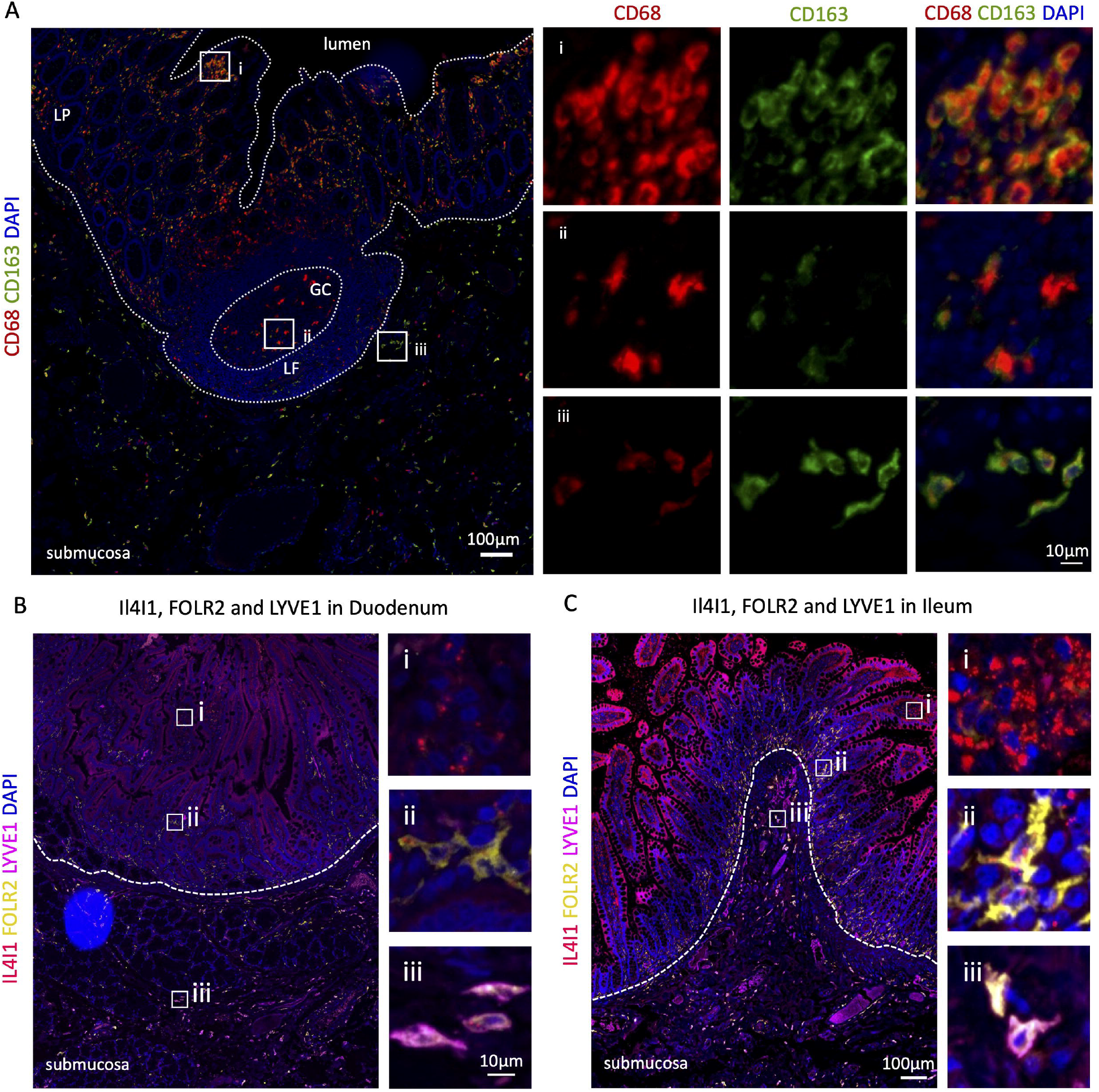
CD68 and CD163 mark spatially distinct macrophage niches in the TME, related to Fig 3. (**A**) Distance quantification workflow shows steps performed to obtain a measurement of macrophage distance to the closest tumor cell from 4-color IF images (**B**) Average protein expression in CD68 Macs and CD163 Macs. (**C-E**) IF images show IL4I1, FOLR2, and panCK signal distribution in (**C**) normal colon mucosa, (**D**) invasive front of CRC, and (**E**) CRC Lymph Node (LN) metastasis. *Left:* Scale bar of 100 μm is identical for all panels on the right. *Right:* Close-up images on the right correspond to boxed regions on the left. The scale bar of 10 μm is identical for all close-up images. DS-desmoplastic stroma, AN-adjacent normal.

**Fig S4.**
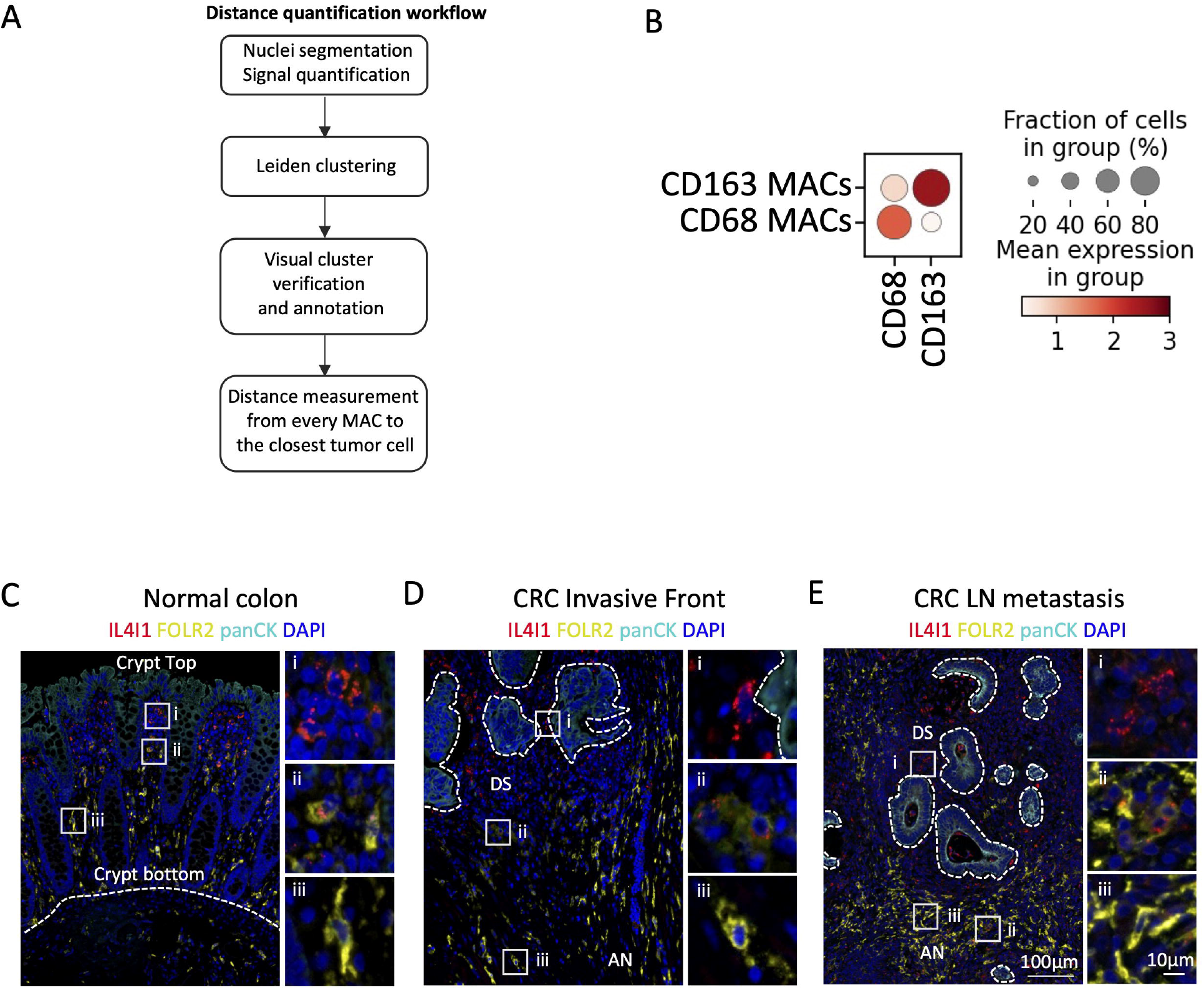
CODEX multichannel imaging reveals cellular interactions in macrophage niches, related to Fig 5. (**A**) Dotplot shows average normalized CODEX marker intensity per identified cell type. (**B**) Dotplot shows average normalized CODEX marker intensity per identified macrophage population. (**C**) Distance (μm) of CD68 TAMs, SPP1 TAMs, CD163 TRMs, FOLR2 TAMs and LYVE1 TRMs to the closest tumor cell. Cells were identified on CODEX images. *P values* were calculated with a linear mixed-effect model with Bonferroni’s corrections for multiple comparisons. (**D**) Barplot shows the distribution of CODEX macrophage populations across macrophage niches. (**E**) Barplot shows the distribution of macrophage niches across CODEX imaged tissue regions.

**Fig S5.**
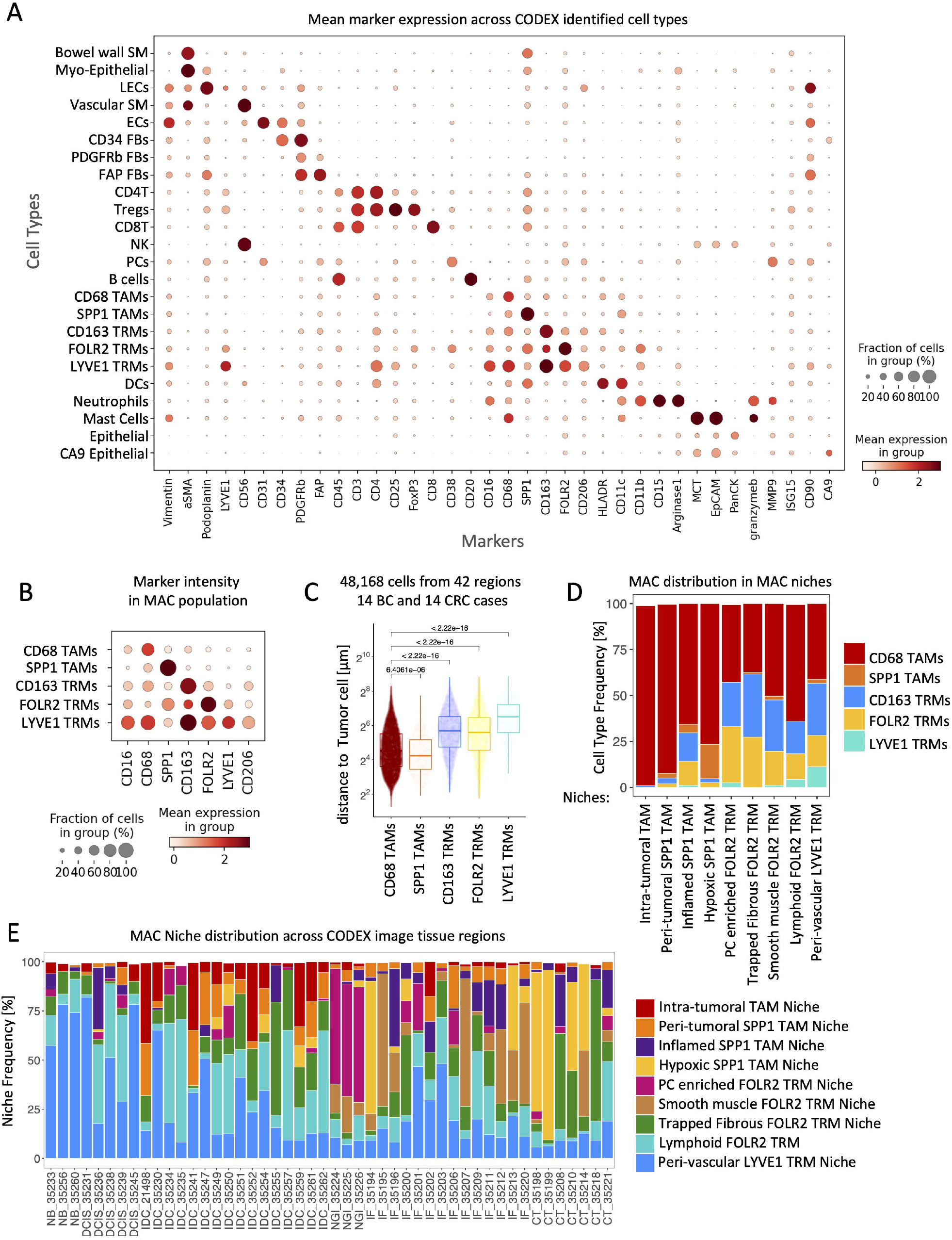
CODEX multichannel imaging, related to Fig 5. (**A**-**I**) Representative *Left:* niche distribution dotplots and *Right:* CODEX images showing cell types enriched in discussed CODEX macrophage niches. Close-up images on the right correspond to boxed regions on the left. Scale bar of 10 μm is identical for all close-up images.

**Fig S6.**
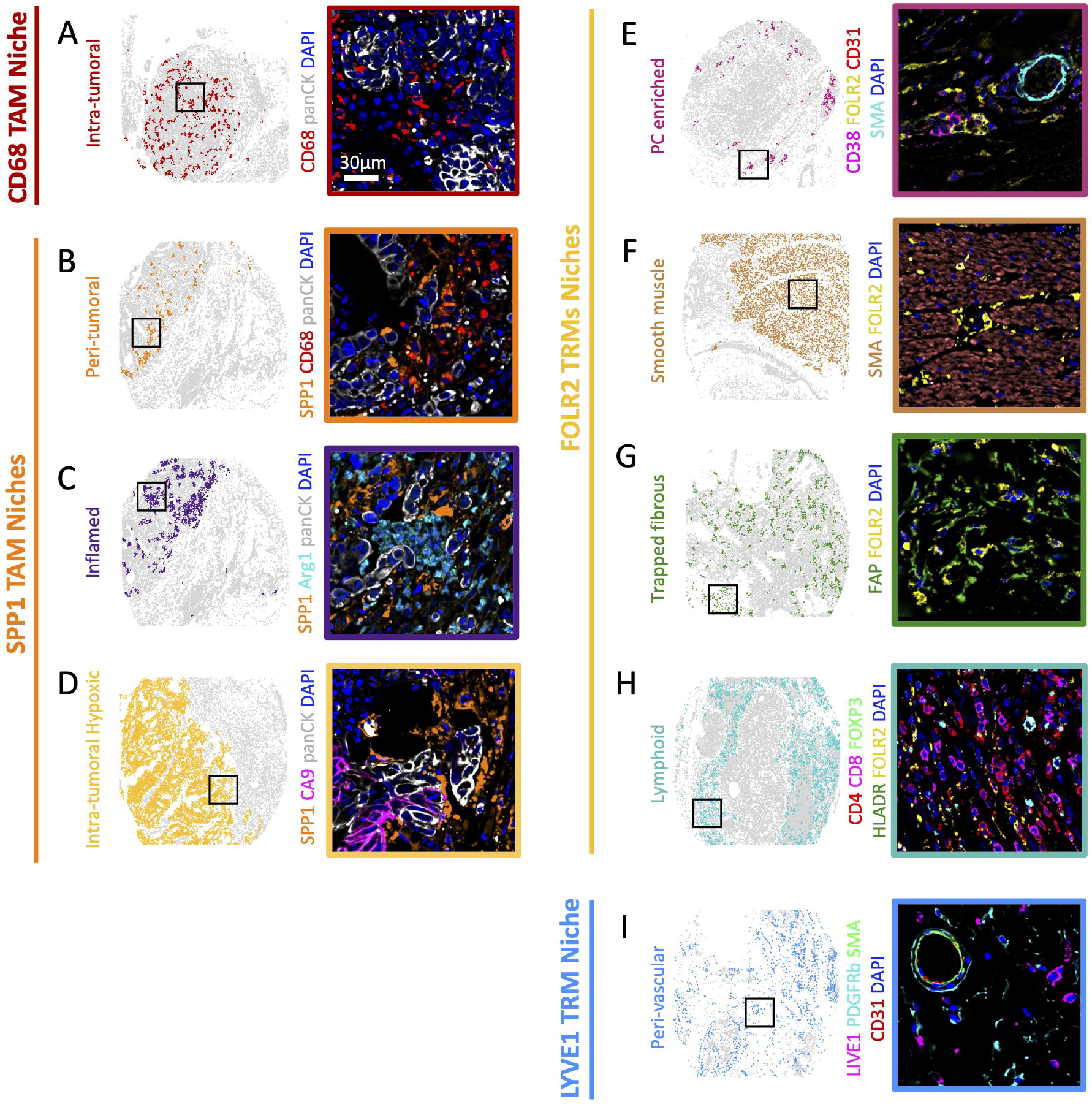
FOLR2 Macrophages and Plasma Cells spatially colocalize, related to Fig 6. (**A**) IF images show PCs marked by CD38 and FOLR2 TRMs marked by FOLR2 in the normal lymph node. Close-up image on the bottom corresponds to boxed regions on the top. (**B**) IF images of PCs marked by co-expression of CD38 and CD138 and FOLR2 TAMs marked by FOLR2 located in normal tissue adjacent to BC. Scale bar of 20 μm is identical for all images. (**C**) Dotplot shows top 10 Ligand and Receptor interactions with the highest communication probability between IgA^+^PCs or IgG^+^PCs and FOLR2 TRMs in CRC scRNA Seq dataset of Lee et. al.

**Fig S7.**
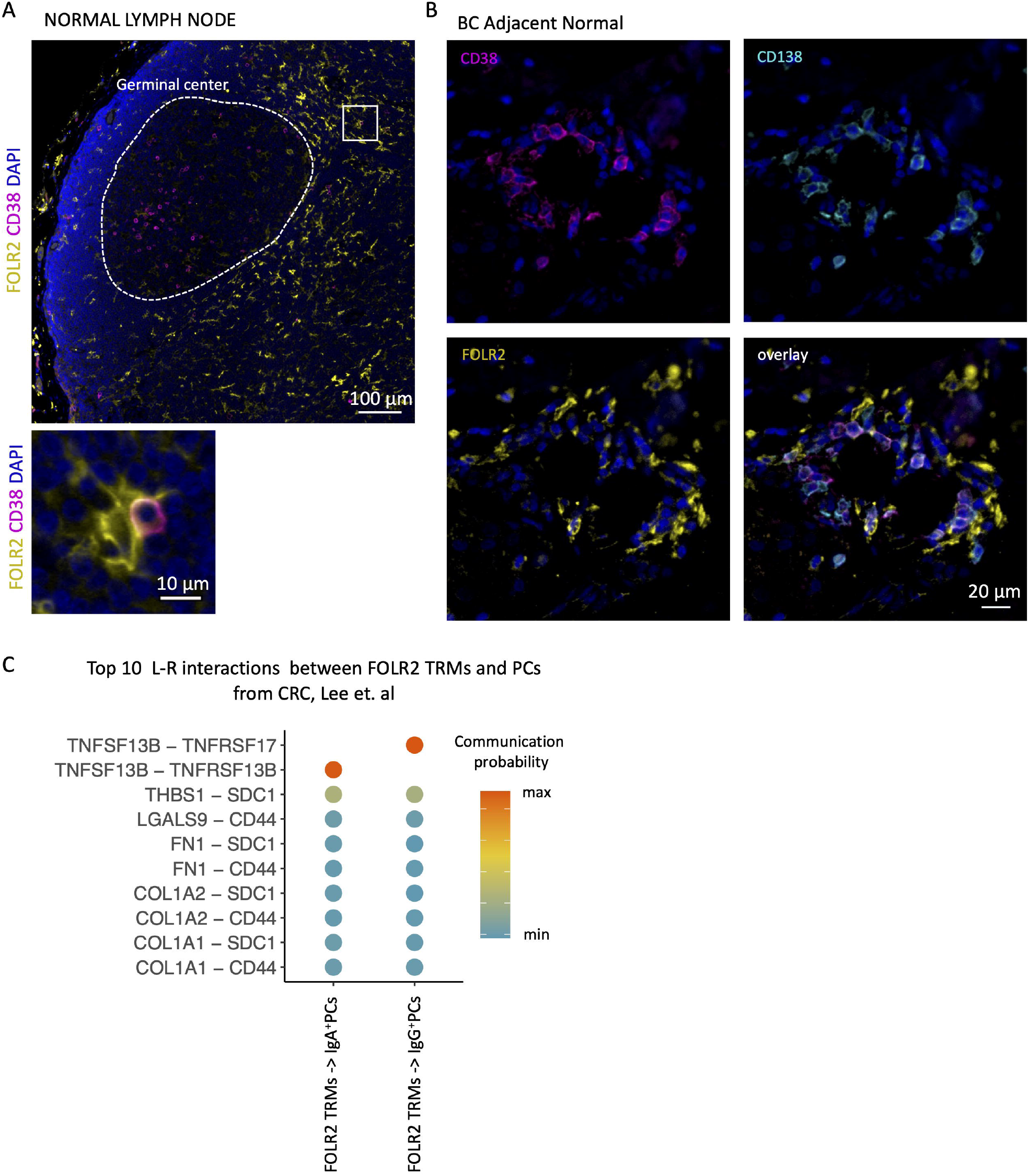
NLRP3 TAMs are associated with neutrophils in Crohn’s Disease, related to Fig 7. (**A**) Dotplot shows the annotation of Tumor (green) and Necrotic (brown) areas (*left*) and normalized expression of SPP1 (*right*) on the 10x Visium FFPE Human Breast Cancer sample. (**B**) Barplot shows normalized log2 SPP1 expression in Tumor and Necrosis regions in **A**. (**C**) Overall survival associations across 16 BC datasets. (**D**) Overall survival associations across 4 CRC datasets.

## Author contributions

M.M., R.W., and M.V.d.R conceived of the study.

M.M., J.W.H., G.L., D.P., G.P.N., B.L. and A.M.N. designed and performed experiments with assistance from S.W.B., S.Z., D.R.C.C.

M.M. and B.L. analyzed the data with assistance from J.W.H., B.L., G.L., L.K., and A.M.N. M.M. and M.V.d.R wrote the paper.

G.C., J.S., R.W., M.V.d.R procured tissue specimens and assisted in data interpretation. All authors commented on the manuscript at all stages.

## Acknowledgments

The authors would like to acknowledge J. Pollack and K.D. Marjon for critical feedback on this work. This work was supported by grants from the National Cancer Institute (R.W., M.V.d.R., R01CA229529), Cancer Research UK (C27165/A29073), and the Virginia and D.K. Ludwig Fund for Cancer Research (M.V.d.R.). J.W.H. was supported by an NIH T32 Fellowship (T32CA196585) and an American Cancer Society—Roaring Fork Valley Postdoctoral Fellowship (PF-20-032-01-CSM). G.L. was supported by Stanford Cancer Institute (SCI) Innovation Award, and by the National Cancer Institute of the National Institutes of Health under Award Number K99CA267171.

## STAR Methods

### RESOURCE AVAILABILITY

#### Lead Contact

Further information and requests for resources should be directed to and will be fulfilled by the Lead Contacts Magdalena Matusiak and Matt van de Rijn.

#### Materials Availability

This study did not generate new unique reagents.

#### Data and Code Availability

CODEX data will be reposited to ImmunoAtlas at https://immunoatlas.org/.

### EXPERIMENTAL MODEL AND SUBJECT DETAILS

#### Human Patient Samples

All clinical specimens in this study were collected with informed consent for research use and were approved by the Stanford University Institutional Review Boards in accordance with the Declaration of Helsinki.

##### Breast and colon cohorts FFPE samples

This study used FFPE samples from 36 invasive breast cancer (IBC) and 32 colon carcinoma (CRC) cases.

##### Crohn’s Disease FFPE samples

We performed the analysis in Fig 7G-H, using three advanced Crohn’s Disease patient FFPE samples.

##### IF and CODEX Tissue microarrays

The tissue microarrays used in ths study were constructed from 36 1.5 mm^2^ regions from 19 CRC cases, and 29 1.5 mm^2^ regions from 18 IBC cases. Regions were selected based on differential spatial staining observed on full section staining with IL4I1, SPP1 and FOLR2 antibodies.

### METHOD DETAILS

#### External datasets

##### Single-cell RNA-seq tumor atlases

We obtained preprocessed scRNA-seq count data from four datasets covering breast carcinoma (BC), and colon carcinoma (CRC). Specifically we used CRC and BC datasets published by Qian et al. (Qian et al. 2020), CRC data from Lee et al. (H.-O. Lee et al. 2020), and BC data from Bassez et al. (Bassez et al. 2021). For each dataset, we extracted monocytes, macrophages, and dendritic cells by clustering SCTransformed count data using Seurat and subsetting clusters expressing AIF1, CST3, CD68, CD163, ITGAX, and HLA-DRA. Next, we integrated the myeloid clusters from the 4 datasets using the reciprocal PCA workflow with Seurat. We used log normalization. To clean the data we excluded dying cells, stressed cells, and cell duplets. We identified dying cells’ clusters by inspecting the distribution of log2(nCount_RNA+1) per cell. Stressed cells were identified based on high expression levels of HSP genes. Cell duplets were identified based on the coexpression of non-myeloid cell markers as follows: myeloid-epithelial cell (TFF3, keratins), myeloid-Tcells (*CD3D*), myeloid-stromal cells (*SPARCL1, SPARC, COL1A1*), and myeloid-plasma (immunoglobulin genes). Since we intended to focus exclusively on monocytes and macrophages, we excluded neutrophils and dendritic cell clusters identified based on the following gene enrichment: neutrophils (SOD2, GOS2, and low detected number of counts per cell), cDC1s (*CLEC9A*), cDC2s (*FCER1A, CD1C, CD1E, and CLEC10A*), migratoryDC (*BIRC3, CCR7, LAMP3*), follicular DC (*FDCSP* and immunoglobulin genes), plasmacytoid DC (*JCHAIN, LILRA4, IRF7*), CD207^+^ DC (*CD1A, CD207, FCAR1A*). Next, we re-clustered the integrated and cleaned Seurat object containing only monocytes and macrophages with resolution = 0.6 in the *FindClusters()* function. We obtained 15 clusters and annotated them based on the most differentially expressed genes in each cluster. Monocytes have been identified by *FN1, FCGR3A*, and *VCAN*. Macrophages were identified based on *C1QA, APOE*, and *TREM2* expression. We merged clusters 0 and 12 into ISG15 TAMs, clusters 1, 6, and 14 into CXCL9 TAMs, and clusters 11 and 13 into LYVE1^+^FOLR2^+^ TRMs. The resulted myeloid object is presented in **Fig 1C**.

##### Spatial transcriptomics

We obtained pre-processed spatial transcriptomic data from Human Breast Cancer: Ductal Carcinoma In Situ, Invasive Carcinoma (FFPE) sample data from 10x website https://www.10xgenomics.com/resources/datasets/human-breast-cancer-ductal-carcinoma-in-situ-invasive-carcinoma-ffpe-1-standard-1-3-0 (**Figure 7 SA-B**).

##### Clinically-annotated tumor transcriptomes

We analyzed 4.231 pre-normalized carcinoma transcriptomes of BC and CRC from the Prediction of Cancer Outcomes using Genomic Profiles (PRECOG) database (Gentles et al. 2015), along with additional datasets listed in Table S4, all of which were processed according to the PRECOG workflow (Gentles et al. 2015). Only datasets with at least 25 samples and available overall survival data were included (Table S4). Specifically, we analyzed 3.905 BC patient samples from 16 datasets and 326 CRC patient samples from 4 datasets.

#### Enrichment of monocyte and macrophage scRNA Seq populations

For the analysis in **Fig 1E, S1C-F**, we selected samples with more than 35 monocyte and macrophage cells and computed the frequency of the different scRNA subsets in each sample. Fig S1C-E, we present these frequencies stratified by tumor type and anatomical location. In addition, for Fig 1E and S1F, we computed a mean frequency for every scRNA subset and calculated a ratio of its frequency between BC and CRC (**Fig 1E**) and normal colon and CRC (**Fig S1F**).

#### Average cluster gene expression

The average gene expression dotplots per scRNA monocyte and macrophage clusters in Fig1D, 2C, 3A, 4H, 7D were plotted using the aggregated myeloid object from Fig 1C.

#### Spatial transcriptomics dataset processing and visualization

For the analysis in **Fig S7A-B**, we used STutility r package to normalize, annotate and visualize the pre-processed spatial transcriptomic data. Specifically, we used the SCTransform function for normalization and the ManualAnnotation function to annotate data based on the H&E image.

#### Immunohistochemistry

For the analysis in Fig 6A and 7A-B, 4 µm tissue sections were deparaffinized and rehydrated. Subsequently, antigen retrieval was performed in EDTA pH 9 buffer for 5 min at 95 °C in a pressure cooker. Slides were next stained with FOLR2, SPP1 or NLRP3 antibodies listed in Table S1, and imaged with a Keyence BZ-X800 microscope at 20′ magnification.

#### Immunofluorescence (IF)

For the analyses shown in Fig 1H, 2A-B, 2E, S2A-C, 3C-E, S3C-E, 5A-C, 6B-D, S6A-B, 7E-H 4µm full tissue sections were deparaffinized and rehydrated. Antigen retrieval was performed using EDTA pH 9 buffer at 95 °C for 10 min. Sections were blocked for 20 min with horse serum and stained for 1h with primary antibodies. Sections were subsequently stained with secondary antibodies for 1 h. A list of primary and secondary antibodies used in this work can be found in Table S1. Sections were then mounted in ProLong Gold Antifade reagent with DAPI and cover-slipped. Stained sections were imaged with a Keyence BZ-X800 microscope at 20′ or 40’ magnification.

#### IF images dearraing

IF images were acquired with a Keyence BZ-X800 microscope at 20′ magnification. Next, the TMA core coordinates were extracted using the dearray functionality in QuPath (Bankhead et al. 2017). Subsequently, the TIFF TMA images were dearrayed using QuPath extracted core coordinates with vips crop function in Linux command line.

#### IF images cell segmentation and immunofluorescence signal quantification

Cell nuclei on the dearrayed TMA cores were segmented using Mesmer (Greenwald et al. 2022). Subsequently, IF signal was quantified for each detected nuclei by computing staining intensity within 3-pixel distance from the nuclear border. We consider a nucleus and its accompanying IF signal within 3-pixel distance from the nuclear border as a cell. In effect, each cell is described by its x and y pixel coordinate and IF staining intensity.

#### Clustering and annotation of IF data

Each individual IF staining was clustered separately. First, IF staining intensity was z normalized using zscore function from scipy.stats python module. Next, cells were clustered using Leiden clustering implementation in scanpy python package. All clusters were individually visually inspected on the dearrayed TIFF images by indicating location of cells attributed to a given cluster. Cell clusters were annotated based on morphology, location, and staining intensity.

For Fig 6C-D, we clustered and annotated cells form 7 1.5 mm^2^ tissue regions including 6 BC and 1 CRC cases. We used FOLR2, IL4I1, and CD138 staining intensity to discriminate FOLR2 TRMs, IL4I1 TAMs and PCs, respectively.

#### Distance quantification of IF and CODEX data

For every TMA core, the distance between every cell and every other cell present in the core was computed using cdist function from scipy.spatial.distance python module. Next, for every macrophage, the shortest distance to a Tumor Cell was selected from the matrix of all cell distances. This shortest distance is reported as the distance to the closest Tumor Cell. For CODEX data, normal breast and gastrointestinal tract samples were excluded.

##### Significance assessment within one tissue region

Wilcox test was used to assess the significance in Figures 3B-E.

##### Significance assessment across multiple tissue regions

Linear mixed-effect models were used to assess significance in Figures 3F-G, 4C, S4C, 6D. We used the lmer function from package lme4 (v1.1.21), and took the tissue region intercept as a random effect. The pairwise p-values were derived from t-ratio statistics in the contrast analysis using the lmerTest (v3.1.2) and corrected for multiple hypothesis testing using the Holm Bonferroni method implemented in the modelbased (v0.1.2) package (github.com/easystats/modelbased).

#### CODEX macrophage distance quantification by niche

For distance quantification in Fig 4C, macrophages were stratified by the macrophage niche they belong to.

#### CODEX antibody panel

The antibody panel in this study was constructed by selecting antibodies targeting epithelial and stromal tumor compartments, with a focus on the myeloid compartment. Detailed information on the included antibodies can be found in Table S2. Each antibody was first conjugated to a unique oligonucleotide tag. Next, antibody-oligonucleotide conjugates were tested in low-plex fluorescence assay to determine whether their staining patterns match patterns established in IHC and IF experiments and to establish the best staining concentration and exposure time. Subsequently, all antibody conjugates were tested together in a single test CODEX imaging multicycle to evaluate optimal concentration, exposure time, and imaging cycle.

#### CODEX imaging

CODEX imaging was performed as previously described (Black et al. 2021). BC and CRC tissue microarrays were simultaneously stained with a previously validated cocktail of antibody-oligonucleotide conjugates and sequentially subjected to CODEX multiplexed imaging using the optimized conditions established during the test run. Metadata with detailed information on each CODEX run can be found in Table S3.

#### CODEX data processing

CODEX imaging data was processed using a software tool called RAPID (Lu G, et al. Manuscript under review, 2022), which included 3D GPU-based deconvolution, spatial drift correction, image stitching, and background subtraction (available at https://github.com/nolanlab/RAPID). Next, cell nuclei segmentation on the processed images was performed using a neural network-based segmentation algorithm called CellVisionSegmenter. CellVisionSegmenter has been shown to work well with segmenting both dense and diffuse cellular tissues with CODEX data (M. Y. Lee et al. 2022). CellVisionSegmenter is an open-source, pre-trained nucleus segmentation and signal quantification software based on the Mask region-convolutional neural network (R-CNN) architecture. The only parameter that was altered was the growth pixels of the nuclear mask, which we found experimentally to work best at a value of 3.

#### CODEX data clustering, visualization, and cell type assignment

Cell clustering and annotation were performed according to a previously published protocol (Hickey et al. 2021). First, nucleated cells were selected by subsetting cells with positive Hoechst signal imaged in 2 separate CODEX cycles. Next, marker signal intensity was z-normalized, and data was overclustered using Leiden clustering in scanpy Python package. Each cluster was visually examined by mapping a location of cells attributed to a given cluster to processed CODEX images and inspecting its marker staining. ImageJ was used to view processed CODEX images. Cell clusters were annotated based on cell morphology, tissue location, and marker staining intensity.

#### CODEX niche analysis

Niche analysis was performed as described earlier by Schurch et al. (Schürch et al. 2020) with k = 10 nearest neighbors and 30 clusters. The cell clusters were annotated and grouped into 13 Niches based on location in the tissue and cell type enrichment score.

#### Ligand-Receptor interaction analysis

Ligan-Receptor analysis was performed using CellChat R package workflow with default settings and using netVisual_bubble function to extract all identified significant liganr-receptor interactions between FOLR2 TRMs and Plasma Cells (PCs). For the analysis in Figure S6D, IgA^+^ and IgG^+^ PCs annotation was extracted from Lee et al. (H.-O. Lee et al. 2020). For Figure 6E, PCs were identified using FindClusters Seurat function with res = 0.4, and selecting cluster #19 with high CD38 and JCHAIN expression. Figure 6E shows all detected significant interactions between FOLR2 TRMs and PCs. Figure S6D shows 10 top significant interactions detected between FOLR2 TRMs and IgA^+^ and IgG^+^ PCs.

#### Gene Set Enrichment Analysis

KEGG pathway gene set enrichment analysis from Figure 5D was performed using clusterProfiler R package. The KEGG enrichment was performed on the list of differentially enriched genes between the 11 transcriptional MAC scRNA Seq populations. Next, enrichment results of Antigen processing and presentation, Phagosome, Lysosome, and Endocytosis gene sets were plotted to compare enrichment of phagocytosis-related pathways between the scRNA MAC populations.

#### Pembrolizumab response analysis

For the analysis in **Fig 4I-J**, was performed on scRNA myeloid transcriptomes form Bassez et al., that we subseted from the aggregated myeloid object form Fig 1D. The patient samples were stratified by the authors of the oryginal publication based on whether the T cell repertoire, as assessed by TCR sequencing, expanded (E) or not (NE) after the pembrolizumab administration. We labeled patients with expanded T cell repertoire as responders (R) and patients with non-expanded T cell repertoire as non-responders (NR).

For the analysis in Fig 4J, we used scRNA monocyte and macrophage transcriptomes of responders and non-responders pre pembrolizumab treatment. We first computed scRNA cluster frequencies in patients with more than 35 monocyte and macrophage cells. Next we compared the mean scRNA cluster frequencies with Chi-squared test using chisq.test function from stats R package and used chisq.posthoc.test function from chisq.posthoc.test R package to asses significance. p values were adjusting using Bonferroni correction.

#### Neutrophil infiltration quantification in BC, CRC, and Crohn’s Disease

For the analysis in Fig 7F, we counted the number of neutrophils present in 1.5 mm^2^ tissue microarray (TMA) cores. The IF-stained TMA cores were evaluated by a pathologist and stratified into cores containing CD68 positive macrophages with diffuse NLRP3 staining or cores that contained CD68 positive macrophages with NLRP3 aggregated in a speck. Cores that contained both diffused and aggregated NLRP3 were classified as cores with NLRP3 speck, as we assumed that the NLRP3 aggregation contains active inflammasome complex that projects the inflammatory signaling. For the analysis in Fig 7H, we counted the number of neutrophils in 1mm^2^ tissue regions selected from whole slide sections. We selected areas that contained CD68 positive macrophages containing NLRP3 aggregated in a speck. Since we didn’t detect any macrophages with NLRP3 diffused staining in the Crohn’s disease tissue sections, we compared the neutrophil numbers in Crohn’s disease patients to benign colon submucosa. CD68 and NLRP3 signals were used to identify NLRP3 TAMs, and Calprotectin was used to identify neutrophils.

#### Survival analyses

For the analyses in Figures 7J and S7C-D, we applied univariable Cox proportional hazards regression to link the relative enrichment of each gene signature (Table S5) to overall survival (survival R package v2.42.3 (Therneau and Grambsch, 2000)) and integrated the resulting z-scores across datasets of the same tumor type as described in (Luca et al. 2021). All survival z-scores were converted to two-sided –log10 p values for clarity.

### QUANTIFICATION AND STATISTICAL ANALYSIS

Wilcoxon test was applied for group comparisons. Linear mixed effect models were applied when groups contained multiple observations from the same tissue region (for instance, when comparing the distance of macrophages to tumor cells across multiple tissue regions). Results with *P* < 0.05 were considered significant. Data analyses were performed with R and python. The investigators were not blinded to allocation during experiments and outcome assessment. No sample-size estimates were performed to ensure adequate power to detect a pre-specified effect size.

---

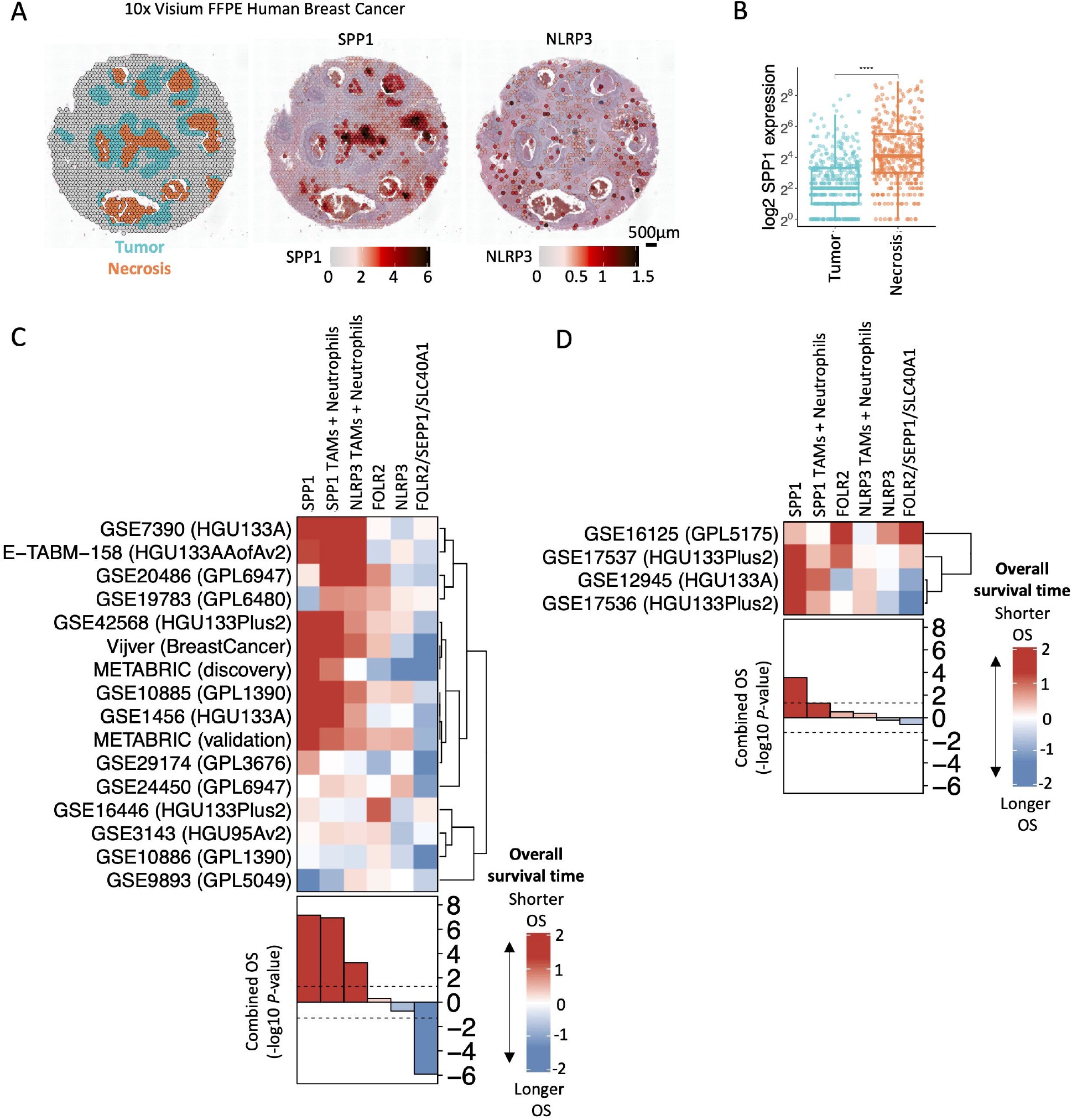

## Notes

### Competing Interest Statement

The authors have declared no competing interest.

### Summary of Updates

Add methods section.

## References

Aguzzi, Adriano, Jan Kranich, and Nike Julia Krautler. 2014. “Follicular Dendritic Cells: Origin, Phenotype, and Function in Health and Disease.” Trends in Immunology 35 (3): 105–13.

Azizi, Elham, Ambrose J. Carr, George Plitas, Andrew E. Cornish, Catherine Konopacki, Sandhya Prabhakaran, Juozas Nainys, et al. 2018. “Single-Cell Map of Diverse Immune Phenotypes in the Breast Tumor Microenvironment.” Cell 174 (5): 1293–1308.e36.

Bassez, Ayse, Hanne Vos, Laurien Van Dyck, Giuseppe Floris, Ingrid Arijs, Christine Desmedt, Bram Boeckx, et al. 2021. “A Single-Cell Map of Intratumoral Changes during Anti-PD1 Treatment of Patients with Breast Cancer.” Nature Medicine 27 (5): 820–32.

Beck, Andrew H., Inigo Espinosa, Badreddin Edris, Rui Li, Kelli Montgomery, Shirley Zhu, Sushama Varma, Robert J. Marinelli, Matt van de Rijn, and Robert B. West. 2009. “The Macrophage Colony-Stimulating Factor 1 Response Signature in Breast Carcinoma.” Clinical Cancer Research: An Official Journal of the American Association for Cancer Research 15 (3): 778–87.

Benson, Micah J., Stacey R. Dillon, Emanuela Castigli, Raif S. Geha, Shengli Xu, Kong-Peng Lam, and Randolph J. Noelle. 2008. “Cutting Edge: The Dependence of Plasma Cells and Independence of Memory B Cells on BAFF and APRIL.” Journal of Immunology 180 (6): 3655–59.

Black, Sarah, Darci Phillips, John W. Hickey, Julia Kennedy-Darling, Vishal G. Venkataraaman, Nikolay Samusik, Yury Goltsev, Christian M. Schürch, and Garry P. Nolan. 2021. “CODEX Multiplexed Tissue Imaging with DNA-Conjugated Antibodies.” Nature Protocols 16 (8): 3802–35.

Chakarov, Svetoslav, Hwee Ying Lim, Leonard Tan, Sheau Yng Lim, Peter See, Josephine Lum, Xiao-Meng Zhang, et al. 2019. “Two Distinct Interstitial Macrophage Populations Coexist across Tissues in Specific Subtissular Niches.” Science 363 (6432). https://doi.org/10.1126/science.aau0964.

Chen, Lih-Chyang, Li-Jie Wang, Nang-Ming Tsang, David M. Ojcius, Chia-Chun Chen, Chun-Nan Ouyang, Chuen Hsueh, et al. 2012. “Tumour Inflammasome-Derived IL-1β Recruits Neutrophils and Improves Local Recurrence-Free Survival in EBV-Induced Nasopharyngeal Carcinoma.” EMBO Molecular Medicine 4 (12): 1276–93.

Coll, Rebecca C., Avril A. B. Robertson, Jae Jin Chae, Sarah C. Higgins, Raúl Muñoz-Planillo, Marco C. Inserra, Irina Vetter, et al. 2015. “A Small-Molecule Inhibitor of the NLRP3 Inflammasome for the Treatment of Inflammatory Diseases.” Nature Medicine 21 (3): 248–55.

Donk, Niels W. C. J. van de, Maarten L. Janmaat, Tuna Mutis, Jeroen J. Lammerts van Bueren, Tahamtan Ahmadi, A. Kate Sasser, Henk M. Lokhorst, and Paul W. H. I. Parren. 2016. “Monoclonal Antibodies Targeting CD38 in Hematological Malignancies and beyond.” Immunological Reviews 270 (1): 95–112.

Fridman, Wolf H., Laurence Zitvogel, Catherine Sautès-Fridman, and Guido Kroemer. 2017. “The Immune Contexture in Cancer Prognosis and Treatment.” Nature Reviews. Clinical Oncology 14 (12): 717–34.

Gentles, Andrew J., Aaron M. Newman, Chih Long Liu, Scott V. Bratman, Weiguo Feng, Dongkyoon Kim, Viswam S. Nair, et al. 2015. “The Prognostic Landscape of Genes and Infiltrating Immune Cells across Human Cancers.” Nature Medicine 21 (8): 938–45.

Goltsev, Yury, Nikolay Samusik, Julia Kennedy-Darling, Salil Bhate, Matthew Hale, Gustavo Vazquez, Sarah Black, and Garry P. Nolan. 2018. “Deep Profiling of Mouse Splenic Architecture with CODEX Multiplexed Imaging.” Cell 174 (4): 968–81.e15.

Gordon, Sydney R., Roy L. Maute, Ben W. Dulken, Gregor Hutter, Benson M. George, Melissa N. McCracken, Rohit Gupta, et al. 2017. “PD-1 Expression by Tumour-Associated Macrophages Inhibits Phagocytosis and Tumour Immunity.” Nature 545 (7655): 495–99.

Gotur, Suhasini Palakshappa, and Vijay Wadhwan. 2020. “Tingible Body Macrophages.” Journal of Oral and Maxillofacial Pathology: JOMFP 24 (3): 418–20.

Hickey, John W., Winston R. Becker, Stephanie A. Nevins, Aaron Horning, Almudena Espin Perez, Roxanne Chiu, Derek C. Chen, et al. 2021. “High Resolution Single Cell Maps Reveals Distinct Cell Organization and Function Across Different Regions of the Human Intestine.” bioRxiv. https://doi.org/10.1101/2021.11.25.469203.

Hickey, John W., Yuqi Tan, Garry P. Nolan, and Yury Goltsev. 2021. “Strategies for Accurate Cell Type Identification in CODEX Multiplexed Imaging Data.” Frontiers in Immunology 12 (August): 727626.

Jiang, Sizun, Chi Ngai Chan, Xavier Rovira-Clavé, Han Chen, Yunhao Bai, Bokai Zhu, Erin McCaffrey, et al. 2022. “Combined Protein and Nucleic Acid Imaging Reveals Virus-Dependent B Cell and Macrophage Immunosuppression of Tissue Microenvironments.” Immunity 55 (6): 1118–34.e8.

Kawakami, Takahiro, Ichiro Mizushima, Kazunori Yamada, Hiroshi Fujii, Kiyoaki Ito, Tetsuhiko Yasuno, Shozo Izui, Masakazu Yamagishi, Bertrand Huard, and Mitsuhiro Kawano. 2019. “Abundant a Proliferation-Inducing Ligand (APRIL)-Producing Macrophages Contribute to Plasma Cell Accumulation in Immunoglobulin G4-Related Disease.” Nephrology, Dialysis, Transplantation: Official Publication of the European Dialysis and Transplant Association - European Renal Association 34 (6): 960–69.

Kennedy-Darling, Julia, Salil S. Bhate, John W. Hickey, Sarah Black, Graham L. Barlow, Gustavo Vazquez, Vishal G. Venkataraaman, et al. 2021. “Highly Multiplexed Tissue Imaging Using Repeated Oligonucleotide Exchange Reaction.” European Journal of Immunology 51 (5): 1262–77.

La Fleur, Linnéa, Vanessa F. Boura, Andrey Alexeyenko, Anders Berglund, Victor Pontén, Johanna S. M. Mattsson, Dijana Djureinovic, et al. 2018. “Expression of Scavenger Receptor MARCO Defines a Targetable Tumor-Associated Macrophage Subset in Non-Small Cell Lung Cancer.” International Journal of Cancer. Journal International Du Cancer 143 (7): 1741–52.

Lee, Hae-Ock, Yourae Hong, Hakki Emre Etlioglu, Yong Beom Cho, Valentina Pomella, Ben Van den Bosch, Jasper Vanhecke, et al. 2020. “Lineage-Dependent Gene Expression Programs Influence the Immune Landscape of Colorectal Cancer.” Nature Genetics 52 (6): 594–603.

Luca, Bogdan A., Chloé B. Steen, Magdalena Matusiak, Armon Azizi, Sushama Varma, Chunfang Zhu, Joanna Przybyl, et al. 2021. “Atlas of Clinically Distinct Cell States and Ecosystems across Human Solid Tumors.” Cell 184 (21): 5482–96.e28.

Mason, James M., Mamta D. Naidu, Michele Barcia, Debra Porti, Sangeeta S. Chavan, and Charles C. Chu. 2004. “IL-4-Induced Gene-1 Is a Leukocyte L-Amino Acid Oxidase with an Unusual Acidic pH Preference and Lysosomal Localization.” Journal of Immunology 173 (7): 4561–67.

Miller, Lloyd S., Eric M. Pietras, Lawrence H. Uricchio, Kathleen Hirano, Shyam Rao, Heping Lin, Ryan M. O’Connell, et al. 2007. “Inflammasome-Mediated Production of IL-1beta Is Required for Neutrophil Recruitment against Staphylococcus Aureus in Vivo.” Journal of Immunology 179 (10): 6933–42.

Mulder, Kevin, Amit Ashok Patel, Wan Ting Kong, Cécile Piot, Evelyn Halitzki, Garett Dunsmore, Shabnam Khalilnezhad, et al. 2021. “Cross-Tissue Single-Cell Landscape of Human Monocytes and Macrophages in Health and Disease.” Immunity 54 (8): 1883–1900.e5.

Nagashima, R., K. Maeda, Y. Imai, and T. Takahashi. 1996. “Lamina Propria Macrophages in the Human Gastrointestinal Mucosa: Their Distribution, Immunohistological Phenotype, and Function.” The Journal of Histochemistry and Cytochemistry: Official Journal of the Histochemistry Society 44 (7): 721–31.

Ramos, Rodrigo, Yoann Missolo-Koussou, Yohan Gerber-Ferder, Christian P. Bromley, Mattia Bugatti, Nicolas Gonzalo Núñez, Jimena Tosello Boari, et al. 2022. “Tissue-Resident FOLR2+ Macrophages Associate with CD8+ T Cell Infiltration in Human Breast Cancer.” Cell 185 (7): 1189–1207.e25.

Okabe, Yasutaka, and Ruslan Medzhitov. 2016. “Tissue Biology Perspective on Macrophages.” Nature Immunology 17 (1): 9–17.

Phillips, Darci, Magdalena Matusiak, Belén Rivero Gutierrez, Salil S. Bhate, Graham L. Barlow, Sizun Jiang, Janos Demeter, et al. 2021. “Immune Cell Topography Predicts Response to PD-1 Blockade in Cutaneous T Cell Lymphoma.” Nature Communications 12 (1): 6726.

Qian, Junbin, Siel Olbrecht, Bram Boeckx, Hanne Vos, Damya Laoui, Emre Etlioglu, Els Wauters, et al. 2020. “A Pan-Cancer Blueprint of the Heterogeneous Tumor Microenvironment Revealed by Single-Cell Profiling.” Cell Research 30 (9): 745–62.

Remmerie, Anneleen, Liesbet Martens, Tinne Thoné, Angela Castoldi, Ruth Seurinck, Benjamin Pavie, Joris Roels, et al. 2020. “Osteopontin Expression Identifies a Subset of Recruited Macrophages Distinct from Kupffer Cells in the Fatty Liver.” Immunity 53 (3): 641–57.e14.

Ridker, Paul M., Jean G. MacFadyen, Tom Thuren, Brendan M. Everett, Peter Libby, Robert J. Glynn, and CANTOS Trial Group. 2017. “Effect of Interleukin-1β Inhibition with Canakinumab on Incident Lung Cancer in Patients with Atherosclerosis: Exploratory Results from a Randomised, Double-Blind, Placebo-Controlled Trial.” The Lancet 390 (10105): 1833–42.

Rőszer, Tamás. 2015. “Understanding the Mysterious M2 Macrophage through Activation Markers and Effector Mechanisms.” Mediators of Inflammation 2015 (May). https://doi.org/10.1155/2015/816460.

Rozanski, Cheryl H., Ramon Arens, Louise M. Carlson, Jayakumar Nair, Lawrence H. Boise, Asher A. Chanan-Khan, Stephen P. Schoenberger, and Kelvin P. Lee. 2011. “Sustained Antibody Responses Depend on CD28 Function in Bone Marrow-Resident Plasma Cells.” The Journal of Experimental Medicine 208 (7): 1435–46.

Schack, Lotte, Romualdas Stapulionis, Brian Christensen, Emil Kofod-Olsen, Uffe B. Skov Sørensen, Thomas Vorup-Jensen, Esben S. Sørensen, and Per Höllsberg. 2009. “Osteopontin Enhances Phagocytosis through a Novel Osteopontin Receptor, the alphaXbeta2 Integrin.” Journal of Immunology 182 (11): 6943–50.

Schürch Christian M., Salil S. Bhate, Graham L. Barlow, Darci J. Phillips, Luca Noti, Inti Zlobec, Pauline Chu, et al. 2020. “Coordinated Cellular Neighborhoods Orchestrate Antitumoral Immunity at the Colorectal Cancer Invasive Front.” Cell 182 (5): 1341–59.e19.

Sharma, Bhesh Raj, and Thirumala-Devi Kanneganti. 2021. “NLRP3 Inflammasome in Cancer and Metabolic Diseases.” Nature Immunology 22 (5): 550–59.

Shin, Yoo-Jin, Hong Lim Kim, Jeong-Sun Choi, Jae-Youn Choi, Jung-Ho Cha, and Mun-Yong Lee. 2011. “Osteopontin: Correlation with Phagocytosis by Brain Macrophages in a Rat Model of Stroke.” Glia 59 (3): 413–23.

Villani, Alexandra-Chloé, Mathieu Lemire, Geneviève Fortin, Edouard Louis, Mark S. Silverberg, Catherine Collette, Nobuyasu Baba, et al. 2009. “Common Variants in the NLRP3 Region Contribute to Crohn’s Disease Susceptibility.” Nature Genetics 41 (1): 71–76.

Xu, Wei, Hyemee Joo, Sandra Clayton, Melissa Dullaers, Marie-Cecile Herve, Derek Blankenship, Maria Teresa De La Morena, et al. 2012. “Macrophages Induce Differentiation of Plasma Cells through CXCL10/IP-10.” The Journal of Experimental Medicine 209 (10): 1813–23, S1–2.

Zhang, Lei, Ziyi Li, Katarzyna M. Skrzypczynska, Qiao Fang, Wei Zhang, Sarah A. O’Brien, Yao He, et al. 2020. “Single-Cell Analyses Inform Mechanisms of Myeloid-Targeted Therapies in Colon Cancer.” Cell 181 (2): 442–59.e29.

## References

Bankhead, Peter, Maurice B. Loughrey, José A. Fernández, Yvonne Dombrowski, Darragh G. McArt, Philip D. Dunne, Stephen McQuaid, et al. 2017. “QuPath: Open Source Software for Digital Pathology Image Analysis.” Scientific Reports 7 (1): 16878.

Greenwald, Noah F., Geneva Miller, Erick Moen, Alex Kong, Adam Kagel, Thomas Dougherty, Christine Camacho Fullaway, et al. 2022. “Whole-Cell Segmentation of Tissue Images with Human-Level Performance Using Large-Scale Data Annotation and Deep Learning.” Nature Biotechnology 40 (4): 555–65.

Lee, Michael Y., Jacob S. Bedia, Salil S. Bhate, Graham L. Barlow, Darci Phillips, Wendy J. Fantl, Garry P. Nolan, and Christian M. Schürch. 2022. “CellSeg: A Robust, Pre-Trained Nucleus Segmentation and Pixel Quantification Software for Highly Multiplexed Fluorescence Images.” BMC Bioinformatics 23 (1): 46.

Schürch, Christian M., Salil S. Bhate, Graham L. Barlow, Darci J. Phillips, Luca Noti, Inti Zlobec, Pauline Chu, et al. 2020. “Coordinated Cellular Neighborhoods Orchestrate Antitumoral Immunity at the Colorectal Cancer Invasive Front.” Cell 182 (5): 1341–59.e19.

